# Shared and genus-specific impact of domestication on mammal oral microbiomes revealed through museomics

**DOI:** 10.64898/2026.10.12.757897

**Authors:** Markella Moraitou, Fawn Young, John L. Richards, Konstantina Saliari, Emmanuel Gilissen, Zena Timmons, Andrew C. Kitchener, Katerina Guschanski

**Affiliations:** Institute of Ecology and Evolution, School of Biological Sciences, University of Edinburgh, Edinburgh, United Kingdom; Natural History Museum Vienna, Vienna, Austria; Department of African Zoology, Royal Museum for Central Africa, Tervuren, Belgium; ULB Center for Diabetes Research, Université Libre de Bruxelles, Brussels, Belgium; Department of Natural Sciences, National Museums Scotland, Edinburgh, United Kingdom; School of Geosciences, University of Edinburgh, Edinburgh, United Kingdom; Department of Ecology and Genetics/Animal Ecology, Uppsala University, Uppsala, Sweden

**Keywords:** museum specimens, dental calculus, metagenomics, pigs, horses, sheep

## Abstract

Animal domestication involves genetic, behavioural and ecological changes, all of which also impact the host-associated microbial communities. Early studies reported diversity loss in the gut microbiomes of domestic and captive animals, but comparisons across a wider diversity of host species showed that loss of microbiota does not always occur. For microbial communities other than the gut, comparisons of multiple domestic-wild pairs are lacking. Here, we use dental calculus metagenomics to investigate the impact of a domesticated lifestyle on the oral microbiome of three pairs of wild and domestic mammals from the genera *Equus* (plains zebra, horse)*, Ovis* (argali, sheep) and *Sus* (wild boar, pig). We find that the oral microbiome of the three domesticates differs both in terms of taxonomic composition and functional capacity from their wild counterparts, but the change is smaller in *Sus* than the other two pairs. While the microbiomes of sheep and horses bear increased resemblance to those of humans compared to their wild relatives, this is unlikely to be due to colonisation by human-derived microbes, as our results suggest that ancestral lineages are mostly retained. Overall, the oral microbiome shows both a consistent signal of domestication, found across all three clades, and a host genus-dependent effect, which likely results from specific changes that underpinned each domestication event.

## Introduction

Domestication, as an evolutionary process, induces morphological, behavioural and physiological changes that enhance adaptation to a novel ecological niche, specifically one defined by co-existence with humans (Purugganan, 2022). As Charles Darwin first observed, many of these changes are common across domestic species (Darwin, 1868; Wilkins et al., 2014), suggesting similar evolutionary pressures under domestication. In mammals, the traits that characterise this ‘domestication syndrome’ (coined by Hammer, 1984) include, among others, docility, smaller teeth, smaller brain or cranial capacity, and juvenile behaviour (Wilkins et al., 2014).

With accumulating discoveries on the importance of microbiomes for host fitness and rapid adaptation under anthropogenic impact (Alberdi et al., 2016; Suzuki & Ley, 2020; Voolstra et al., 2026), studies have investigated the impact of domestication (Bae et al., 2025; Metcalf et al., 2017; Prabhu et al., 2020; Reese et al., 2021) and captivity on the gut microbiome (Clayton et al., 2016; Fatoretto et al., 2022; McKenzie et al., 2017; Metcalf et al., 2017; Nishida & Ochman, 2021; Pimiento & Antonelli, 2022). An early study on primates suggested captivity leads to considerable loss of diversity and convergence towards a more human-like gut microbiome (Clayton et al., 2016). This ‘humanisation’ of the gut microbiome has been replicated in captive primates and seems at least partly driven by the displacement of ancestral microbes by human-derived strains (Nishida & Ochman, 2021). However, studies across a wider diversity of mammals find that the impact of captivity on microbiome composition and diversity is not always the same, may vary across hosts (McKenzie et al., 2017; Narat et al., 2020; Prabhu et al., 2020) and the specific circumstance of captivity, such as altered diet (often a simpler diet with less fibre), habitat (often cleaner and less diverse), stress and antibiotic use (Dallas & Warne, 2023; Kuthyar et al., 2023). Specifically, primates and carnivorans show the most drastic gut microbiome changes, while ruminant artiodactyls tend to be rather stable both in terms of composition and microbiome richness (McKenzie et al., 2017). Against expectations, rhinoceros microbiomes were found to increase in diversity in captivity (McKenzie et al., 2017), although this finding was not replicated by a later study (Gibson et al., 2019). Overall, despite the effects of captivity, the gut microbiome retains a strong signature of host taxonomy, e.g., at the genus level (McKenzie et al., 2017). Similarly, while domestication affects the gut microbiome, the signature of host phylogeny persists (Reese et al., 2021). The changes in the animal gut during domestication are at least partly due to dietary changes, often reflecting increased digestible carbohydrate and reduced fibre intake (Reese et al., 2021), but can also be independent of diet (Prabhu et al., 2020). As seems to have occurred in the human gut during industrialisation (Sanders et al., 2023), domestication can lead to the decrease or loss of beneficial symbionts (Prabhu et al., 2020), but can also result in increased diversity depending on the specific ecological changes that come with it, e.g., whether the post-domestication diet is less or more varied (Reese et al., 2021).

While most current knowledge on the effects of domestication and captivity on animal microbiomes is based on the gut microbiome, the oral microbiome is also likely to be involved. Like the gut microbiome, the oral microbiome is shaped by host diet and habitat (Moraitou et al., 2026), but at the same time is subject to different dynamics than the gut. Specifically, the oral microbiome tends to be more conserved across hosts (Asangba et al., 2022) and through time (Farese et al., 2026; Gancz et al., 2023), and more exposed to environmental microbiota (Shaiber et al., 2020). Therefore, findings cannot be extrapolated between the two body sites without investigation. Particularly relevant in the context of domestication, the oral microbiome has adapted to an increase in starch consumption, as manifested in the increased abundance of amylase-binding streptococci during the course of human evolution (encoding for proteins that sequester salivary α-amylases; Fellows Yates et al., 2021). It is known that some domestic and commensal animals, such as dogs, pigs and mice, due to their proximity to and association with humans, have adapted to an increased consumption of starch via duplications of the amylase gene (Pajic et al., 2019). However, it is not known if their oral microbiomes have also adapted in a similar way to those of humans.

Here, we use dental calculus metagenomes to investigate the effect of domestication on the oral microbiome of mammals. We focus specifically on three pairs of wild/domestic animals: the domestic pig (*Sus domesticus*) and the Eurasian wild pig (*Sus scrofa*, which we will refer to as the wild boar for clarity); the domestic horse (*Equus caballus*) and the plains zebra (*Equus quagga*); and, finally, the domestic sheep (*Ovis aries*) and the argali (*Ovis ammon*). The latter two pairs do not include the wild progenitors of the domestic species, the Przewalski’s horse (*E. przewalskii*; Gaunitz et al., 2018) and the Asian moufflon (*O. gmelini*; Her et al., 2022), respectively, but instead include a close wild relative in the same genus. *Ovis* and *Sus* were both domesticated approximately 11,000 years before present during the Neolithic (Vigne et al., 2009; Zeder, 2008), whereas *Equus* was domesticated much later, approximately 4,000 years before present (Librado et al., 2021). Our aim is to investigate if there is a consistent effect of domestication on oral microbiome composition, diversity and function across the three wild-domestic pairs. More specifically, we explore if these changes reflect a ‘humanisation’ of the oral microbiome by analysing them alongside publicly available human dental calculus metagenomes. To investigate the potential effect of diet on the functional capacity of the oral microbiome, we focus on genes related to starch metabolism, specifically microbial amylases, as well as the amylase-binding protein abpA, a surface protein encoded by some oral *Streptococcus* species to bind and utilise host salivary amylases. Finally, we use metagenome-assembled genomes (MAGs) to understand if ancestral microbiome lineages are retained following domestication or if they are replaced by human microbiota.

## Materials and Methods

### Dental calculus metagenomic dataset

The dataset consisted of 107 dental calculus metagenomes from six non-human mammal species (Moraitou et al., 2025, 2026), representing three genera with one wild and one domesticated counterpart, as well as 14 human dental calculus metagenome samples, nine from historical individuals from Fellows Yates et al. (2021) and five from modern individuals from Velsko et al. (2019). The non-human mammal species were chosen to represent the closest domestic-wild pairs for which data were available: the domestic sheep (*Ovis aries*) and the argali (*Ovis ammon*), the domestic pig (*Sus domesticus*) and the wild boar (*Sus scrofa*), and the domestic horse (*Equus caballus*) and the plains zebra (*Equus quagga*). Note that 24 of the 29 *Ovis aries* are feral, belonging to the St. Kilda Soay sheep population in Scotland (Boyd et al., 1964). For some analyses these were treated as a distinct host group, referred to as “feral” and separate from the fully domestic sheep (Table 1, Supplementary Table 1). In addition, to control for contamination, we included five environmental control metagenomes (derived from swabs of the museum collection area associated with these specific specimens) and 36 blank samples that were processed alongside these particular dental calculus samples as negative controls for DNA extraction and library preparation.

**Table 1.** Dataset summary. Number of dental calculus metagenomes analysed per species.

| Scientific name | Common name | Domestication status | Host reference genome for mapping | Number of metagenomes |
| --- | --- | --- | --- | --- |
| *Equus caballus* | Domestic horse | Domestic | GCF_002863925.1 | 6 |
| *Equus quagga* | Plains zebra | Wild | GCA_026770645.1 | 24 |
| *Ovis aries* | Domestic sheep | Feral | GCF_016772045.2 | 24 |
|  |  | Domestic | GCF_016772045.2 | 5 |
| *Ovis ammon* | Argali | Wild | GCA_028583565.1 | 10 |
| *Sus domesticus* | Domestic pig | Domestic | GCA_017957985.1 | 12 |
| *Sus scrofa* | Wild boar (or Eurasian wild pig) | Wild | GCF_000003025.6 | 26 |
| *Homo sapiens* | Human | NA | GCA_000001405.29 | 14 (9 historical and 5 modern) |
| Total dental calculus metagenomes |  |  |  | 121 |
| Environmental swabs |  |  |  | 5 |
| Extraction blanks |  |  |  | 18 |
| Library preparation blanks |  |  |  | 18 |
| Total metagenomes |  |  |  | 162 |

### Sequence pre-processing

The sequences were processed as described in Moraitou et al. (2026). Briefly, paired-end reads were merged using fastp (S. Chen et al., 2018). At the same step, any Illumina adaptors and barcodes were trimmed, as were polyX and polyG tails (an artifact of Illumina NovaSeq two-colour chemistry) and low-quality bases (mean quality below 30 using a sliding window of 3bp). Finally, merged reads shorter than 30bp were filtered out. PCR duplicates were removed using the ‘dedupe’ module from BBMap (Bushnell, 2014).

The retained reads were mapped against a concatenated genomic reference including the human genome (GCA_000001405.29), the PhiX virus genome (a common spike-in for Illumina sequencing; GCA_000819615.1), and the genome of the host species (Table 1), using bwa aln with settings ‘-n 0.04 –k 2 –l 1024 –o 2’ (H. Li & Durbin, 2009). For human samples, as well as for blanks and environmental swabs, the mapping was done only against the human and PhiX genomes. Following mapping, only unmapped reads were extracted for downstream analysis using SAMtools (Danecek et al., 2021), as these should predominantly consist of microbiome-derived sequences. To assess if these metagenomic reads indeed represented the oral microbial community, we performed source-tracking using decOM (Duitama González et al., 2023) with a custom source matrix including human oral (N = 177), terrestrial mammal oral (N = 49), marine mammal oral (N = 52), rumen (N = 40), skin (N = 81), and sediment/soil metagenomes (N = 79), as described in Moraitou et al. (2025, 2026).

### Taxonomic and Functional Profiling

To taxonomically and functionally characterise the dental calculus metagenomes, we used a contig-based approach as described in Moraitou et al. (2026). In summary, metagenomic reads were assembled into contigs using MEGAHIT (D. Li et al., 2016), separately for each sample. Only contigs > 1000bp were retained and combined across all samples in a single file, which was then dereplicated at 98% nucleotide identity using the MMseqs2 easy-cluster function (Hauser et al., 2016) with settings ‘--min-seq-id 0.98 –c 0.8 –-cov-mode 5’. Contigs were taxonomically classified using the Contig Annotation Tool (CAT) (von Meijenfeldt et al., 2019) against a reformatted version of the GTDB database (Parks et al., 2022). Functional annotations were performed using DRAM (Shaffer et al., 2020), using 1000bp as the minimum contig length to include all retained contigs.

To obtain taxon and gene abundances, we mapped the metagenomic reads from dental calculus and control samples back to the combined and dereplicated contig file using the KMA mapper (Clausen et al., 2018) with settings ‘-nc –1t1’ and extracted the number of unique reads per sample mapping to each contig. For the taxon abundances, we considered reads mapping with at least 98% average sequence identity to contigs that were classified at least to the family level and which were at least 10% covered by reads (breadth of coverage). For gene abundances, we used similar criteria, requiring that at least 50% of each contig was covered by metagenomic reads.

### Binning and Evaluation of Metagenome-Assembled Genomes

MAGs were assembled using the nf-core/mag pipeline (Krakau et al., 2022) with the ‘--ancient_dna’ module enabled. Their completeness and contamination was estimated with CheckM (Parks et al., 2015), and those of medium quality and above (completeness ≥ 50% and contamination ≤ 10%) according to MIMAG (minimum information about a metagenome-assembled genome; Bowers et al., 2017) were taxonomically classified and used to construct de-novo phylogenies with GTDBtk (Chaumeil et al., 2022). To quantify the abundance of these MAGs in each sample, we first obtained a dereplicated set of MAGs using the ‘dereplicate’ function from dRep at 98% sequence identity (Olm et al., 2017) and then mapped the metagenomic reads against this dereplicated set using the KMA mapper. We considered a MAG present in a sample if it had > 10,000 mapped reads, breadth of coverage > 75%, and average nucleotide identity > 98% (Moraitou et al., 2026).

### In-silico decontamination

The taxon and gene tables were analysed alongside the sample metadata using the phyloseq R package (McMurdie & Holmes, 2013). For the taxon table, only classifications assigned to at least the species level were retained, and any subspecies classifications were agglomerated (‘tax_glom’ function). The taxon abundances of the negative controls were then used to identify contaminants in our dataset. We used a similar approach as presented in Moraitou et al. (2026), by removing any taxa that had a lower average relative abundance in dental calculus than in controls. Specifically, we first calculated the average relative abundance within host species, and then across host species, to account for the varying number of samples per species. If this value was lower than the average relative abundance in controls (extraction blanks, library blanks, environmental swabs), the taxon was considered a contaminant and removed. In addition, we removed taxa that were found in less than 20% of the samples of any single host species. Finally, relative abundances <0.01% in any given sample were set to zero to reduce spurious identifications.

Dental calculus samples with less than 10,000 reads classified to the species level were excluded from downstream analyses. Those that showed a higher proportion of skin and soil content than of oral microbial communities (based on the decOM source-tracking results) were considered poorly preserved and were also removed.

For the gene table, any entries assigned to species identified as contaminants were removed, and then the abundances of annotated genes were summed (regardless of the microbial taxon to which they belonged) to get the total abundance of each gene in a given metagenome. Genes with less than 100 reads and samples identified as poorly preserved based on taxonomic analysis or containing less than 1 million total reads (insufficient depth to recover gene content; Supplementary Figure 1) were excluded. To further reduce the size of the dataset and exclude spurious assignments, we only retained genes found in ≥10% of the samples of at least one host species.

### Species richness and taxonomic composition

As variation in read depth can bias estimates of species richness, a rarefaction analysis was performed using the ‘rarefy’ function from the vegan R package (Oksanen et al., 2022). According to the rarefaction curves, richness plateaued at approximately 20,000 reads (Supplementary Figure 2), therefore we rarefied to this depth for comparisons of species richness. We tested for the effects of host genus, domestication status and their interaction using an ANOVA (analysis of variance). Even after rarefying, the number of reads mapping to contigs was included as an explanatory variable in the ANOVA model, to ensure we capture any residual effect of read depth. The normality of residuals was evaluated using a Shapiro-Wilk test to ensure that the assumptions of the model were met. Within-factor comparisons of significant factors were performed using Tukey’s HSD (Honest Significant Difference) tests.

To account for the compositional nature of the microbiome data (Gloor et al., 2017) during community-wide analysis, the abundances of the decontaminated taxon table were transformed using a CLR (centred-log ratio) normalisation, as well as a phylogenetically informed PhiLR normalisation (phylogenetic isometric log ratio; Silverman et al., 2017) based on the GTDB phylogenetic tree. A binary table representing the presence and absence of taxa was also used.

Differences in composition were visualised using a Redundancy Analysis (RDA) implemented in the microViz R package (Barnett et al., 2021) using host species, except for human, as constraints, and tested using PERMANOVA with genus, domestication status and their interaction, as well as the number of reads as predictors. Both RDA and PERMANOVA were run three times for each of the three transformations: binary presence/absence, CLR-transformed abundances and PhILR-transformed abundances. For the PERMANOVA, Jaccard distances were used for presence-absence and Euclidean distances were used for CLR-transformed (also referred to as Aitchison distances; Aitchison, 1986) and PhILR-transformed data. To ensure that there is no multicollinearity between the constraints of the RDA, a stepwise model selection and variance inflation calculations were performed. Finally, multivariate variances (distance of each sample to its host species centroid) were estimated using the ‘betadisper’ function from the vegan R package with Aitchison distances. Between-host group differences were then tested with an ANOVA and Tukey’s HSD.

To investigate if the oral microbiomes of domesticates bear closer resemblance to the human oral microbiome than to those of their wild counterparts, we compared the distances of each non-human sample to the human centroid across different host groups and performed pairwise testing using Kruskal-Wallis tests (adjusting p-values for multiple testing using the Holm method). This, again, was done using Jaccard distances, Aitchison distances (Euclidean distances of CLR-transformed abundances) and Euclidean distances on PhILR-transformed abundances.

### Differential abundance analysis and loss/gain of taxa

Differential abundance analysis between wild and domesticated animals was performed using MCMCglmm (Hadfield, 2010; Sweeny et al., 2023) for the 100 most abundant microbial genera in each host genus, using CLR-normalised abundances as the response variable and allowing for different intercepts and interaction effects for both domestication status and host genus (after excluding humans and feral sheep). No adjustment of pMCMC was performed, as standard frequentist approaches for multiple-testing adjustment may not be appropriate for Bayesian p-values, but instead we used a more stringent threshold of pMCMC < 0.01. To understand if the detected shifts in abundance are consistent across all three wild-domestic pairs, we also ran the analysis separately for each of the three host genera, without the host genus interaction term. Finally, to put the above analyses into context we identified which of these taxa are typical of human oral microbiota by testing humans against all wild animals for the same microbial genera. This allowed us to assess whether the changes in abundance between wild and domesticates may reflect a ‘humanisation’ of the oral microbiome as a result of domestication.

Beyond changes in abundance, we aimed to identify taxa that are lost or gained in domesticated animals. To do this, we first detected structural zeros (true presence/absence) across host species using ANCOM-BC2 (Lin & Peddada, 2024), for all genus-level taxa and with host species as the primary variable of interest (again excluding humans and feral sheep). The structural zeros output was then used to identify loss and gain of taxa in domesticates. If a microbial genus was present in the domesticated species and absent in its wild counterpart, it was considered a ‘gain’, whereas the reverse pattern was considered a ‘loss’.

### Functional capacity of the microbiome

Three different approaches were used to investigate the functional capacity of the oral microbiome of domesticates and wild species: gene abundances, KEGG pathway abundances, and functional trait completeness inferred with distillR (Alberdi, 2025). The KEGG pathways were obtained by mapping the KEGG annotations from DRAM (after filtering and decontaminating the gene table) to pathways using the KEGGREST R package (Tenenbaum & Maintainer, 2024). Pathway name and class were retrieved with the ‘keggGet’ function. After compiling lists of all genes in each pathway, the completeness (percentage of genes present) and abundance (as the total gene abundance) of each pathway were calculated for the entire dataset and separately for every sample. Pathways that were unrelated to microbiome function (e.g., referring to human diseases, organismal systems and viruses) or with less than 20% of genes present in the dataset were removed as spurious. Functional traits were identified with the distillR R package (Alberdi, 2025) and further aggregated to the compound level, producing a table with functional annotations such as ‘methionine degradation’ and ‘betaine biosynthesis’ and their completeness in each sample.

Gene and pathway abundances were CLR-normalised before downstream analysis, but functional trait values from distillR were not, because they were quantified in terms of completeness, not abundances. For all three, the compositional differences were visualised using RDA ordinations, the effects of genus, domestication status and their interaction were tested using PERMANOVA, while also accounting for the number of reads. Differential abundance analysis of all retained KEGG pathways was performed as described for genus-level taxonomic abundances.

To investigate potential adaptation of the oral microbiome to starch consumption, we focused on the abundances of genes encoding for microbial amylases and the amylase-binding protein abpA. Amylase genes were identified by filtering the DRAM annotations for enzyme committee (EC) numbers 3.2.1.1 (α-amylases), 3.2.1.2 (β-amylases), 3.2.1.3 (γ-amylases) and 3.2.1.133 (maltogenic amylases). We then tested if their overall relative abundance differed by domestication status (domestic, feral, wild), diet (with *Ovis* and *Equus* denoted as herbivores and *Sus* as omnivores), the domestication-diet interaction, and, finally, read count using ANOVA and Tukey’s HSD. Amylase-binding proteins were not annotated by DRAM, so they were identified manually using HMMER3 (Eddy, 2009). Specifically, we used ‘hmmbuild’ to create an HMM profile for abpA based on an alignment of 33 protein sequences from various *Streptococcus* species as well as five *Gemella* and one *Granulicatella* genomes (Supplementary Table 2). Then, we used ‘hmmsearch’ to search for matching sequences across the translated open reading frames identified by DRAM, and for any matches we extracted mapping and taxonomic information of the contigs in which they were located. The per-sample abundances of any annotations were calculated based on contig mapping statistics, using the same thresholds as described above (98% average nucleotide identity and 50% breadth of coverage). Hits were validated by searching against the RefSeq protein database using protein-protein BLAST (Madden, 2003).

### Phylogenetic analysis of MAGs

To understand if domesticated animals have been colonised by human-derived microorganisms due to proximity with humans, we relied on the bacterial and archaeal phylogenetic trees created using the GTDBtk ‘de_novo_wf’ workflow and pruned to include only MAGs with completeness >75% and contamination <5%.

To test for associations between MAGs found in domesticated animals to those found in their wild counterparts or in humans, two different approaches were used, one based on tree topologies and the other based on phylogenetic distances. For both approaches, we considered all feral sheep as domesticated, with the logic that they have undergone domestication in the past.

In the topology-based analysis we compared the frequency of occurrence of nodes with three different topological patterns: ((domestic, wild), human); ((domestic, human), wild); and ((human, wild), domestic). The ((domestic, wild), human) topology, or ((D,W),H) for short, and the ((D,H),W) topology represent higher relatedness between a MAG isolated from a domesticated animal with a MAG isolated from either its wild counterpart or a human, respectively. The ((H, W), D) topology represents a less likely scenario that groups human-derived MAGs and wild animal MAGs to the exclusion of domesticated MAGs. Cases in which the wild and domesticated animals were not from same wild-domesticated pair were not considered in this analysis. MAG presence was established using both assembly– and mapping-based approaches (see above). To identify these nodes, we first considered all nodes that split MAGs from any single host species on one side, and MAGs from any two host species (exactly) on the other side. Valid nodes with the ((D,W),H), ((D,H),W) and ((H,W),D) topology had to have ‘D’ and ‘W’ MAGs from the same host genus, otherwise they were classified as “other”. To assess the significance of the observed patterns, a permutation test was performed, randomly shuffling host species labels (while retaining MAG identity) 100 times, and the frequencies of different node types were calculated again. The observed counts and proportions of node types were compared to the permuted results. For the distance-based analysis, we focused on MAGs found in domesticates and calculated the phylogenetic distance to the nearest wild counterpart MAG and to the nearest human MAG. As the point of this analysis was to understand if domesticates retain their ancestral MAGs or have them replaced by a related human lineage, we limited its phylogenetic scope by requiring that either the human or the wild MAG comes from the same genus as the domestic one. MAGs that were identified in more than one host species via mapping were represented in the phylogeny as multiple tips (one per host species) with a branch length of zero.

## Results

### Sequence processing

After mapping against the combined human, host and PhiX reference genomes, the dental calculus samples retained a median of 17,442,101 reads (range: 152 – 128,570,126) and the negative controls (including museum environmental swabs, extraction and library preparation blanks) had a median of 828 reads (range: 0 – 670,379) (Supplementary Table 1).

At least one contig could be assembled from 119 of the 121 dental calculus samples (median number of contigs per sample: 35,667, range: 1 – 194,338) and 115 of these had at least one contig longer than 1,000 bp. The median number of contigs > 1,000 bp in these samples was 5,861 (range: 1 – 33,534). Contigs could be assembled from only eight negative controls (range: 1 – 151 contigs per negative control) with only seven contigs longer than 1000bp, all in museum environmental swabs (Supplementary Table 1).

For the contig-based analysis of the metagenomes, we characterised a reference set of 685,589 contigs, which were obtained after clustering all contigs longer than 1000 bp across all samples at 98% identity. In total, we identified 7,779 microbial species-level taxa in our dataset (7,520 bacteria and 259 archaea), whose per-sample abundances were determined by mapping the metagenomic reads to the reference contigs. After decontamination and abundance filtering, 2,641 of these taxa were retained (Supplementary Figure 3, Supplementary Table 3). We also removed eight samples as poorly preserved, either because they had less than 10,000 classified reads or because they had a high skin and soil microbial proportion compared to oral, according to the decOM source-tracking results (Supplementary Figure 4, Supplementary Table 4). After this procedure, we retained 113 samples for downstream analyses.

### Microbial richness varies among domesticated species

We identified a median of 205 microbial species (range: 16 – 414) among 112 samples with at least 20,000 classified reads (Figure 1A, Supplementary Table 5). Host genus was the most important variable explaining species richness (Table 2), with all pairwise comparisons among non-human genera showing significant differences (Figure 1A, Supplementary Table 6). With the exception for *Ovis*-*Homo*, all other comparisons with human were also significant, with *Homo* showing lower oral microbial species richness. Notably, the species of the genus *Sus* (domestic pig and wild boar) had the most diverse microbiomes, which is typical for mammals with omnivorous diets (Moraitou et al., 2026).

**Figure 1.**
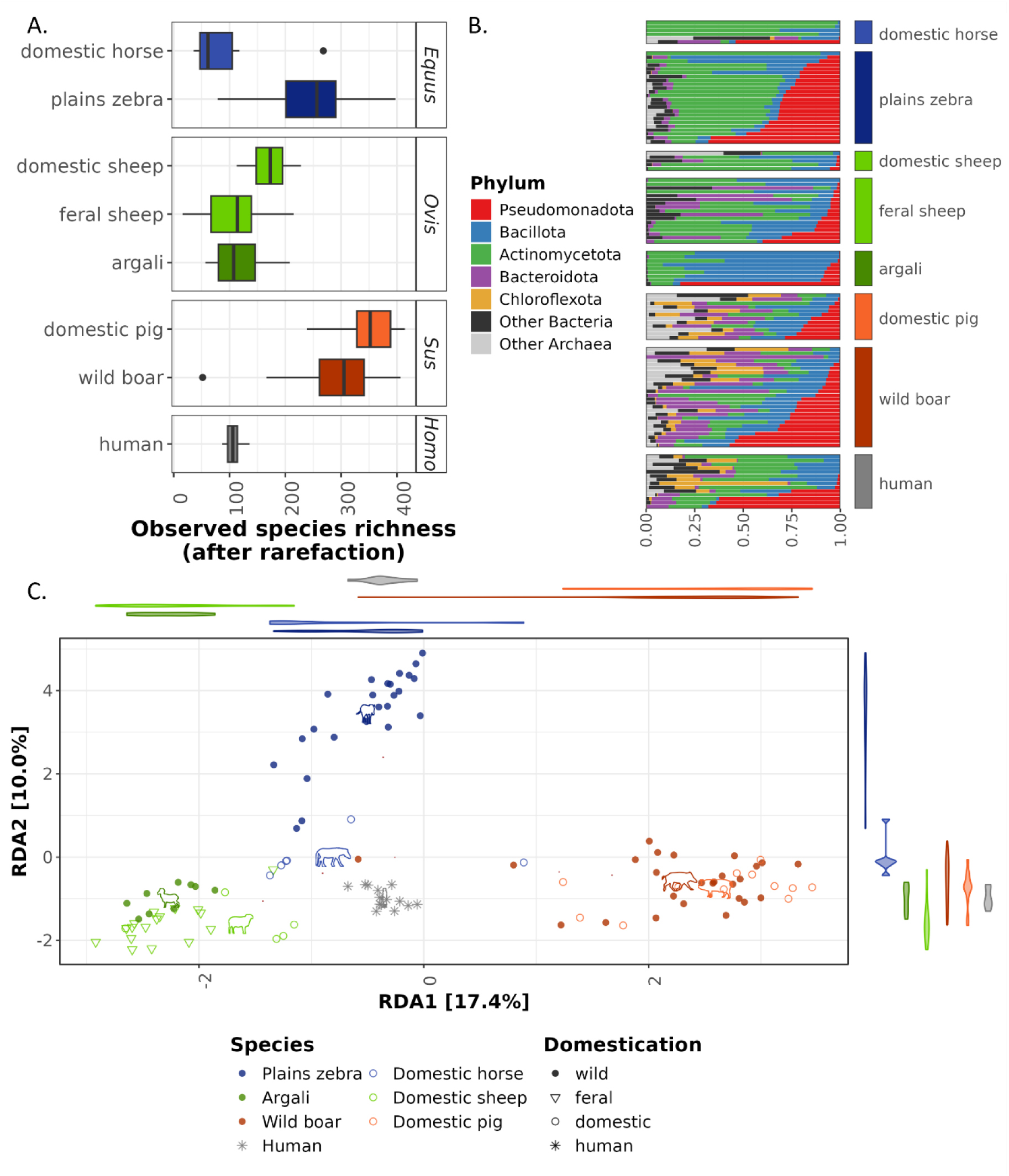
Comparison of taxonomic composition and richness. **A)** Species richness of dental calculus microbiome in domesticated and wild study species and humans, after decontamination and rarefaction to 20,000 classified reads. **B)** Phylum-level microbiome composition across host groups. **C)** Redundancy analysis (RDA) ordination on CLR-transformed abundances, using host species as constraints (except for human). Different colours correspond to host species and genera and different shapes indicate the domestication status (open circles – domesticated, open triangles – feral, closed circles – wild). The violin plots along the axes show the distributions of the host species along that axis. The phylopic icons, plotted using the rphylopic R package (Gearty & Jones, 2023), indicate the centroid of each host species (for sheep the centroid represents only the fully domestic sheep). Note that sheep (*Ovis aries*) samples are split into feral (triangles) and truly domestic sheep (circles) in all panels.

**Table 2.** Effect of host genus and domestication on species richness. ANOVA (analysis of variance) results on species richness (after decontamination and rarefaction) using as explanatory variables host genus, domestication status as well as the interaction between them, after accounting for the number of reads mapping to contigs (to account for the data content of each sample).

| Term | Sum Sq | F-value | p-value |
| --- | --- | --- | --- |
| Contig-mapped reads | 47703.3 | 11.44 | 0.001 |
| Host genus | 716656.8 | 57.29 | < 0.001 |
| Domestication | 6008.8 | 0.72 | 0.489 |
| Host genus : Domestication | 132460.7 | 15.88 | < 0.001 |
| Residuals | 354677.1 |  |  |

Domestication status did not have a consistent directional effect and instead affected richness differently in different host genera (Figure 1A, Table 2). The genus *Equus* showed significantly lower diversity in domestic horses than in plains zebras (Supplementary Table 6, Figure 1A). In contrast, in the other two genera, domesticated animals showed higher richness, albeit not significantly so. The feral sheep of St Kilda and the wild argali both had lower microbiome richness than the domestic sheep (Figure 1A).

### Humanisation of the oral microbiome evident in sheep and horses, but not pigs

Across all host species the oral microbiome was dominated by three bacterial phyla: Pseudomonadota, Bacillota and Actinomycetota (Figure 1B). In addition, in microbiomes of the two omnivorous genera *Sus* (wild boar and domestic pig) and *Homo*, we also detected considerable proportions of the bacterial phylum Chloroflexota, as well as several archaeal phyla (Figure 1B), as is typical of animalivorous and omnivorous mammals (Moraitou et al., 2026).

Host genus was the most important factor explaining oral microbiome composition both in terms of presence/absence and abundance of microbial species (Figure 1C, Supplementary Figure 5A,B) and accounting for 15-29% of variance depending on the distance metric (PERMANOVA p-value < 0.001; Supplementary Table 7). Domestication status, as well as its interaction with host genus, had smaller but still significant effects, explaining 4-5% of the variances for all three metrics (p-value < 0.001; Supplementary Table 7). The number of reads had a small but significant impact across all metrics, but as expected it was larger for Jaccard distances (R^2^ = 3%, p-value = 0.001) compared to CLR– or PhILR-normalised abundances (R^2^ = 1%, p-value = 0.01; Supplementary Table 7).

Human samples clustered near the centre of the first and second ordination axes of the ordination, regardless of the distance metric used (Figure 1C, Supplementary Figure 5A,B). Domestic horses and sheep were shifted towards the centre compared to their wild counterparts, zebras and argalis, respectively, indicating a trend towards the humanisation of the oral microbiome in domesticates. Supporting this interpretation, we found smaller distances between the dental calculus microbiomes of domesticates and the human centroid (Figure 2), but they were realised in a different way in the two genera. Specifically, the horse oral microbiome was more similar to the human centroid when considering abundance-informed metrics (CLR– and PhILR), but not for presence/absence-based distances, whereas the opposite was true for sheep (Figure 2). The feral sheep of St. Kilda showed intermediate levels of similarity to humans, being placed between the wild argali and fully domestic sheep, but closer to the wild animals, in line with alpha diversity results (Figure 1A). In the genus *Sus* we found no evidence of humanisation of the oral microbiome, regardless of metric (Figure 2).

**Figure 2.**
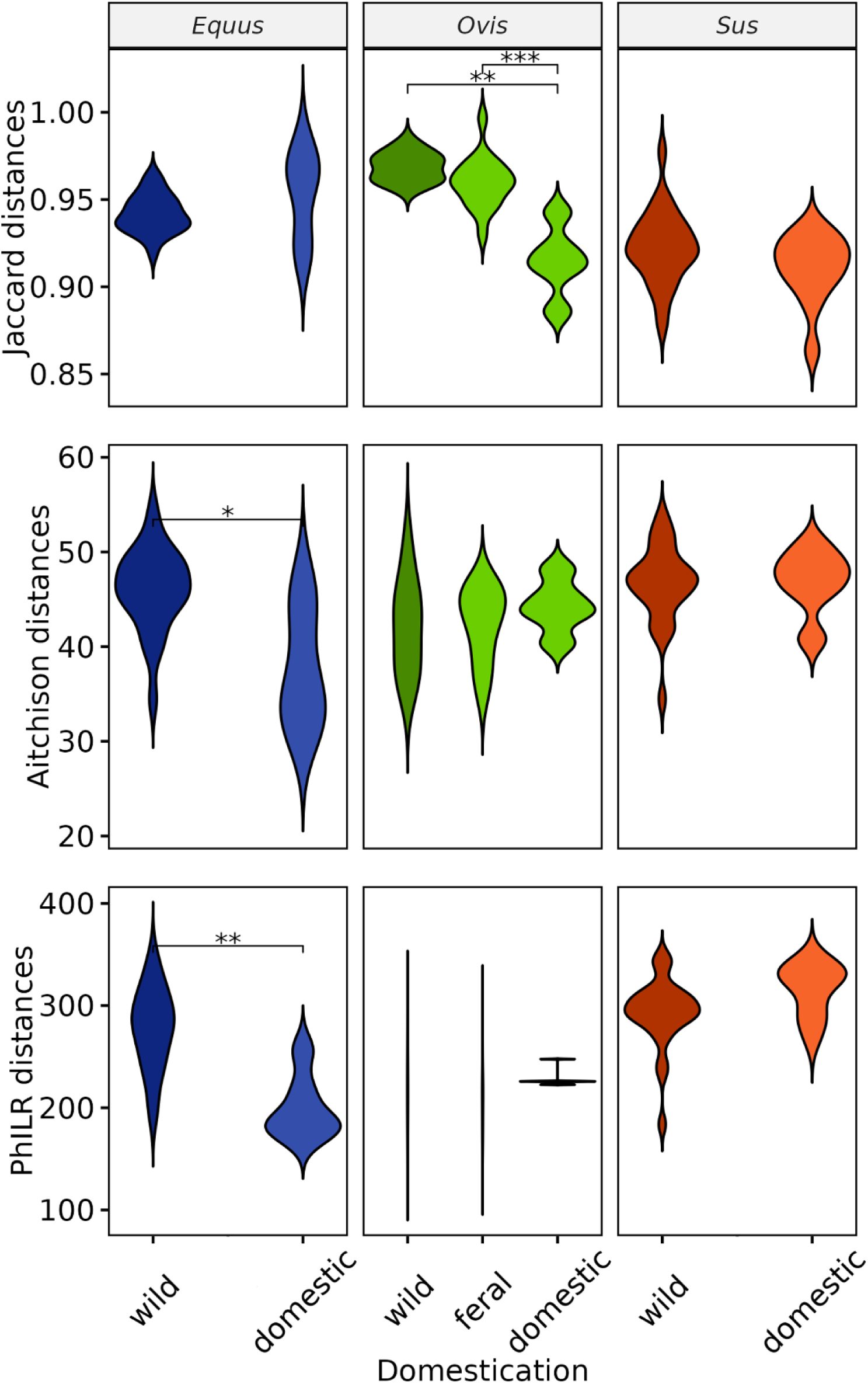
Distance of the taxonomic composition of each sample to the human centroid, calculated using different data normalisations and distance metrics. Top: Jaccard distances on presence-absence data. Middle: Aitchison distances (Euclidean distances on CLR-transformed data). Bottom: Euclidean distances on PhILR-transformed data. Asterisks indicate significant differences between the different domestication categories of a host genus, identified using a Kruskal-Wallis test: “*” for p < 0.05, “**” for p < 0.01, “***” for p < 0.001.

Despite this observed humanisation in domestic *Equus* and *Ovis*, when comparing the multivariate variance (average distance to the group centroid) of wild and domesticated microbiomes (across all genera), we found no significant differences (ANOVA: p = 0.36), suggesting that domesticate microbiomes were not significantly more similar to each other than the wild microbiomes. However, multivariate variance did differ between host groups (ANOVA: F=4.8, p < 0.001). Specifically, wild boars had the most variable microbial communities, significantly so compared to feral sheep, human and argali samples (Tukey’s HSD: p-values < 0.05; Supplementary Table 8; Supplementary Figure 5C).

### Consistent shifts in microbiome abundances present in domesticates

Differential abundance analysis was run using MCMCglmm for 189 genus-level taxa, representing the 100 most abundant microbial genera in each non-human host genus. Across the three host genera we observed repeated, consistent shifts in taxon abundances between wild and domesticated counterparts for several microbial taxa (using pMCMC < 0.01 to identify the most robust results; Supplementary Table 9; Figure 3A). One of these, the typical oral archaeon *Methanobrevibacter* (Granehäll et al., 2021; Pilliol et al., 2024), was enriched in domesticates in all three independent within-genus comparisons. Several others show a significant positive effect in one host genus and a positive trend, albeit not significant, in one or in both remaining host genera, such as *Olsenella_F* (typical oral taxon; Dewhirst et al., 2001) and Anaerovoraceae taxon RUG099 (isolated from the horse gut; Gilroy et al., 2022) (Figure 3A; Supplementary Figure 6). On the other hand, we also found taxa consistently depleted in domestic animals: the most notable example is *Basfia_A*, a taxon isolated from the rumen (Kuhnert et al., 2010), but also reported in the oral cavity. In addition, several more taxa were depleted in domestic horses or sheep with non-significant negative trends in the other genera, e.g., the ruminant-associated *Mannheimia* (Angen et al., 1999) and the oral commensal *Streptococcus* (Abranches et al., 2018) (Figure 3A; Supplementary Figure 6). Overall, we found a larger number of significant comparisons within *Equus* (N = 41) and *Ovis* (N = 26) than within *Sus* (N = 8; Supplementary Table 9). Among the domestic-associated taxa, some were also enriched in humans, most notably *Olsenella_F* and *Methanobrevibacter_A* (Figure 3A; Supplementary Figure 6), providing further evidence for the humanisation of the domesticated oral microbiome, which may occur either due to horizontal transfer of symbionts or due to similar host filtering conditions in humans and domesticates. Conversely, those taxa that are consistently depleted in domesticates also show low abundance in humans compared to wild animals. This is most marked, unsurprisingly, in the ruminant-associated *Basfia*_A and *Mannheimia* (Angen et al., 1999; Kuhnert et al., 2010), but also seen in typical oral taxa, such as *Streptococcus*, *Lautropia* and *Neisseria* (Figure 3A).

**Figure 3.**
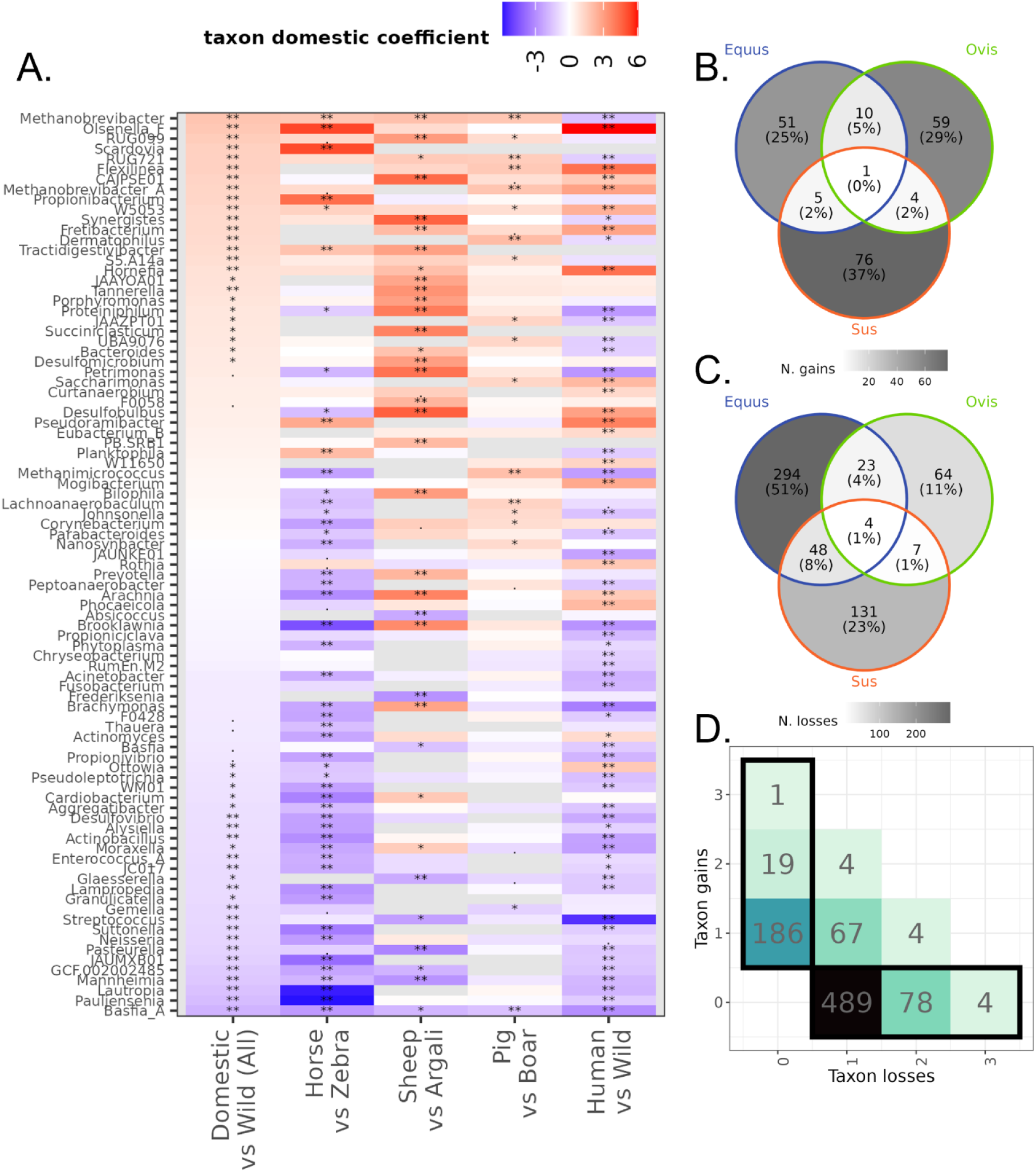
Taxon gains, losses and abundance changes. **A)** Results of four runs of differential abundance analysis with MCMCglmm. From left to right: all domestic against all wild samples (while accounting for host genus, and after removing feral sheep and humans), domestic horses against plains zebras, domestic sheep against argalis, domestic pigs against wild boars, and humans versus all wild species. Only taxa that were differentially abundant in a least one comparison are presented (using pMCMC < 0.01) and ranked according to the estimated coefficient, indicating abundance change in domesticates in the first comparison. Colour signifies the estimated coefficient with domestic species or humans as a reference (red: enriched in domesticates/humans, blue: enriched in wild). Asterisks indicate a comparison that was significant and passed the sensitivity analysis: “***” for pMCMC < 0.001, “**” for pMCMC < 0.01, “*” for pMCMC < 0.05, “.” for pMCMC < 0.1. **B)** Genus-level gains of taxa in domesticates, excluding taxa that are lost in even one of the wild-domestic pairs. **C)** Genus-level losses of taxa in domesticates, excluding taxa that are gained in even one of the wild-domesticated pairs. **D)** Summary of frequency of loss or gain of taxa in domesticates across the three wild-domestic pairs in this study. The numbers indicate the number of genus-level taxa in a given category (for instance, 67 taxa are lost in one wild-domestic pair, but gained in another, while 186 taxa are gained in one pair and never lost). This value is also reflected in the tile colour. The black outlines highlight taxa that are only ever lost (taxon losses > 0 and taxon gains = 0; same taxa as panel **B**) or only ever gained (taxon gains > 0 and taxon losses = 0; same taxa as panel **C**).

### Several taxa are consistently lost or gained in domesticates

Next, we investigated if domestication leads to consistent turnover of specific microbial taxa. Losses were more frequent (N = 736) than gains (N = 306), both overall and within each host genus (Figure 3D, Supplementary Table 10), with most events occurring in a single wild-domestic pair (Figure 3B,C). However, we identified one taxon that was consistently gained in domesticates of all three host genera (Figure 3B,D): *Olsenella*, and more specifically a clade related to *O. uli*, a common member of the human oral microbiome (Dewhirst et al., 2001). Note that the related clade *Olsenella_F* was also present in wild hosts (as are other *Olsenella* clades), but was nonetheless more abundant in domestic species (Figure 3A; Supplementary Table 9).

Four taxa were consistently lost in domesticates (Figure 3C,D): *Enterococcus_A* (a common mammalian gut commensal; Aarestrup et al., 2002), *Simonsiella* (found in the oral cavity of warm-blooded vertebrates; Hedlund & Tønjum, 2015), *Saccharicrinis* (isolated from marine environments; Bachmann, 1955; Yang et al., 2014) and *Psychrobacter* (isolated from seawater and processed meat; Juni & Heym, 1986) (Supplementary Table 10). The first two broadly agree with the differential abundance analysis, as *Enterococcus_A* was significantly depleted in domesticates and *Simonsiella* showed a similar but non-significant trend (Supplementary Table 9). *Saccharicrinis* and *Psychrobacter* were not abundant enough to be included in differential abundance analysis and therefore may reflect spurious classifications or occasional external contamination. Among all domesticates, horses have lost the most taxa (Figure 3C), in agreement with their reduced species richness (Figure 1A). Perhaps unexpectedly, even domestic pigs and sheep showed more losses than gains, despite having a higher species richness than their wild counterparts (Figure 1A). This could be because losses and gains were calculated per host species, whereas alpha diversity was assessed per sample. However, there may also be a technical reason, as the calculation of structural zeros with ANCOMBC required a 10% prevalence cut-off, meaning it is ignoring some rare taxa.

### Domestication may lead to loss of functional potential in the oral microbiome

Functional analysis was performed on the same set of contigs used for taxonomic profiling. We identified 20,912 unique features, 12,630 of which were retained after decontaminating the gene table and filtering out low-prevalence genes. 111 samples were sufficiently preserved for this analysis. Of the retained genes, the 5,829 KEGG annotations accounted for 96% of total gene abundance and mapped to 358 unique KEGG pathways. After filtering spurious annotations and pathways with low completeness, 102 pathways were retained (Supplementary Table 11). We also identified 145 functional traits with distillR, 76 related to compound degradation, 62 related to biosynthesis, and seven related to structural components (Supplementary Table 12).

As observed for microbial taxonomic composition, host evolutionary relationships were also the main drivers of functional composition (Figure 4A, Supplementary Figure 7). Regardless of metric (gene abundances, pathway abundances, or distillR functional trait completeness), host genus explained the largest percentage of variance at 25-27% (p-values = 0.001; Supplementary Table 13). Domestication status, both independently and as an interaction with host genus, had a significant effect in all cases. For gene and pathway abundances, the independent effect (6% and 10% of variance, respectively; p-values < 0.001) was larger than the interaction (about 5% for both; p-values < 0.005; Supplementary Table 13). For the functional trait completeness, the independent effect of domestication (3%, p-value = 0.014), was slightly smaller than the interaction (4%, p-value = 0.004; Supplementary Table 13).

**Figure 4.**
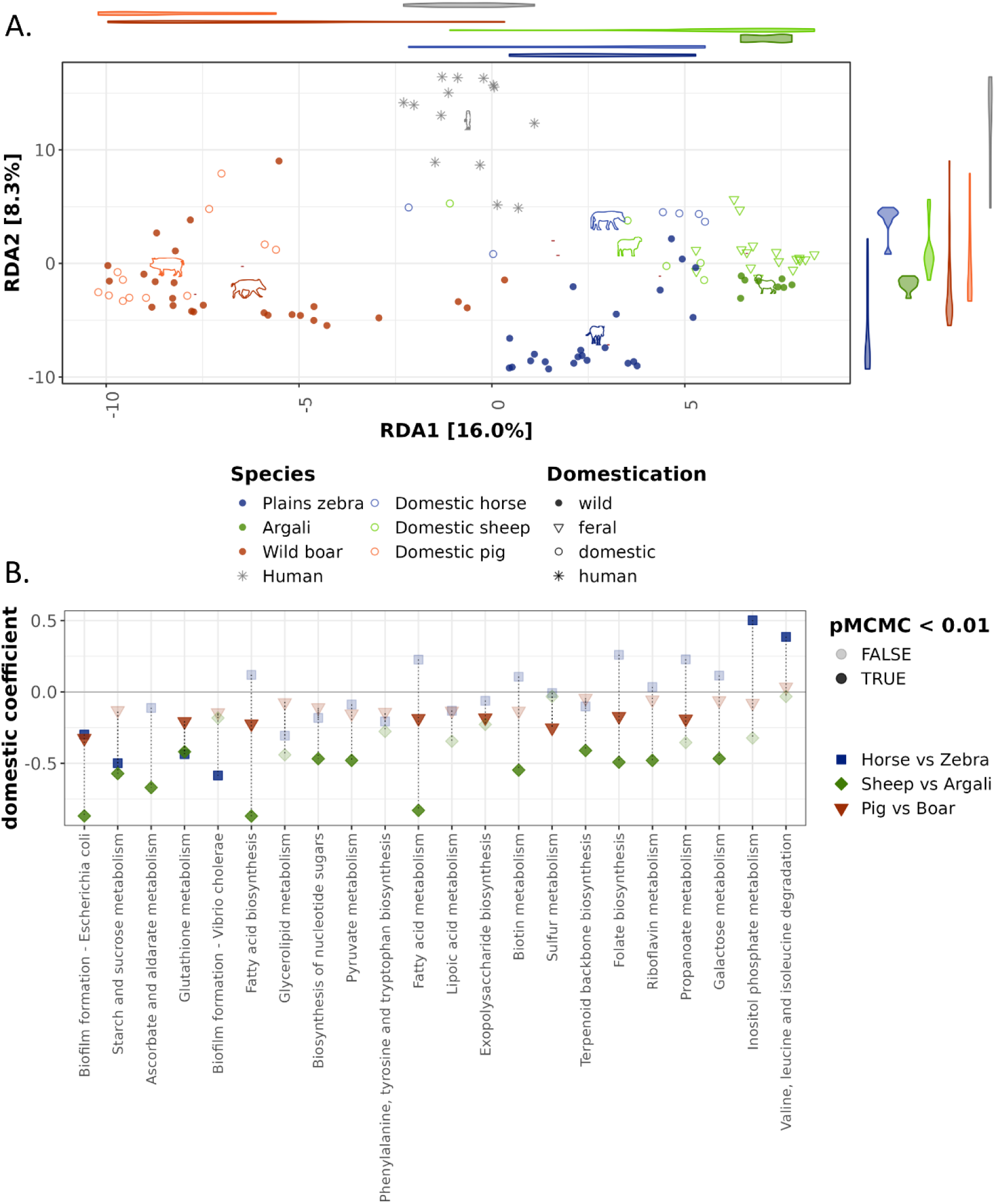
Functional comparison of the oral microbiome. **A)** Redundancy analysis ordination on CLR-transformed gene abundances using host species as constraints. Host species is designated by different colours, whereas the domestication status is shown with different shapes. The violin plots along the axes show the distributions of the host species along that axis. The phylopic icons indicate the centroid of each host species. **B)** Estimated log-fold changes in abundance for the KEGG pathways that were found to differ significantly in at least one comparison: domestic horses compared to plains zebras (blue square), domestic sheep against argalis (green diamond), and domestic pig against wild boar (orange triangle). Higher opacity of the points indicates significant comparisons (MCMCglmm pMCMC < 0.01).

Domestic animal microbiomes were characterised by a depletion of KEGG pathways. Out of 102 KEGG pathways only two were enriched in domestic animals (MCMCglmm pMCMC < 0.01; Supplementary Table 14), and both of them only in the comparison of horses vs. zebras, whereas 17 pathways were significantly depleted in at least one and often two domestic-wild comparisons (most often in sheep and pigs; Figure 4B; Supplementary Table 14). The most consistently depleted pathways were related to polysaccharide metabolism, such as the biosynthesis of poly-N-acetyl-glucosamine (PGA) exopolysaccharide (part of the ‘Biofilm formation – *Escherichia coli*’ pathway) and cellulose degradation by beta-glucosidase (part of the ‘Starch and sucrose metabolism’ pathway) (Figure 4B; Supplementary Table 14). The decreased abundance of the beta-glucosidase gene, a key enzyme for cellulose degradation, in domestic animals may be in response to a more digestible diet with less plant fibre. The reason for the decreased abundance of the PGA gene is less obvious, but it has previously been associated with biofilm formation and increased virulence of periodontal pathogens (Shanmugam et al., 2015).

Despite domesticated microbiomes being more similar to those of humans in terms of taxonomic composition (Figure 2), we obtained mixed evidence for the functional profiles. We found no increased similarity between domesticates and humans in terms of gene abundances, and in terms of pathway abundances domestic sheep were functionally more human-like than feral sheep, whereas, surprisingly wild zebras were more human-like than domestic horses (Supplementary Figure 8).

### Domesticated herbivore microbiomes respond to increased starch consumption

As starch is often added to the feed of domestic animals (Julliand et al., 2006; Lunesu et al., 2021) and is known to impact the microbiome, we specifically focused on the abundances of amylases, the enzymes responsible for digesting starch. We found that α-amylases are the most abundant amylase type (Figure 5), mostly represented by the KEGG orthologue K05343 (maltose alpha-D-glucosyltransferase / alpha-amylase [EC:5.4.99.16 3.2.1.1]). Total relative abundance of amylases was strongly affected by domestication status (ANOVA F = 23.9 p-value < 0.001), while no other factors (diet, diet-domestication interaction and read count) were significant (F < 2, p-value > 0.2). Unexpectedly, the feral sheep microbiomes encoded for most amylase genes, whereas domestic and wild microbiomes did not differ significantly (Table 3). Still, domestic sheep have on average a higher relative abundance of amylases than argalis (0.15% and 0.05%, respectively), as is the case for horses compared to zebras (0.21% and 0.16%). However, this not the case for domestic pigs compared to wild boars, which have comparable values (ca. 0.18% for both) (Figure 5). Microbial amylases are encoded by a diversity of oral taxa such as *Actinomyces, Arachnia*, *Streptococcus, Flexilinea* and *Propionibacterium*. However, in horses and sheep the amylase profiles appear each to be dominated by a single genus: *Propionibacterium* in domestic horses and *Actinomyces* in domestic and feral sheep (Figure 5). This mirrors the differences in abundance of these taxa, as *Actinomyces* was most abundant in feral sheep and *Propionibacterium* in domestic horses (Supplementary Figure 6), possibly indicating that these taxa increased in abundance in response to the diet shift towards starch-containing food items. Consistent with this, the amylase profiles of wild boars and domestic pigs were more similar to each other, which is expected given that wild boars are omnivores, and therefore their diet may not have changed considerably with domestication.

**Figure 5.**
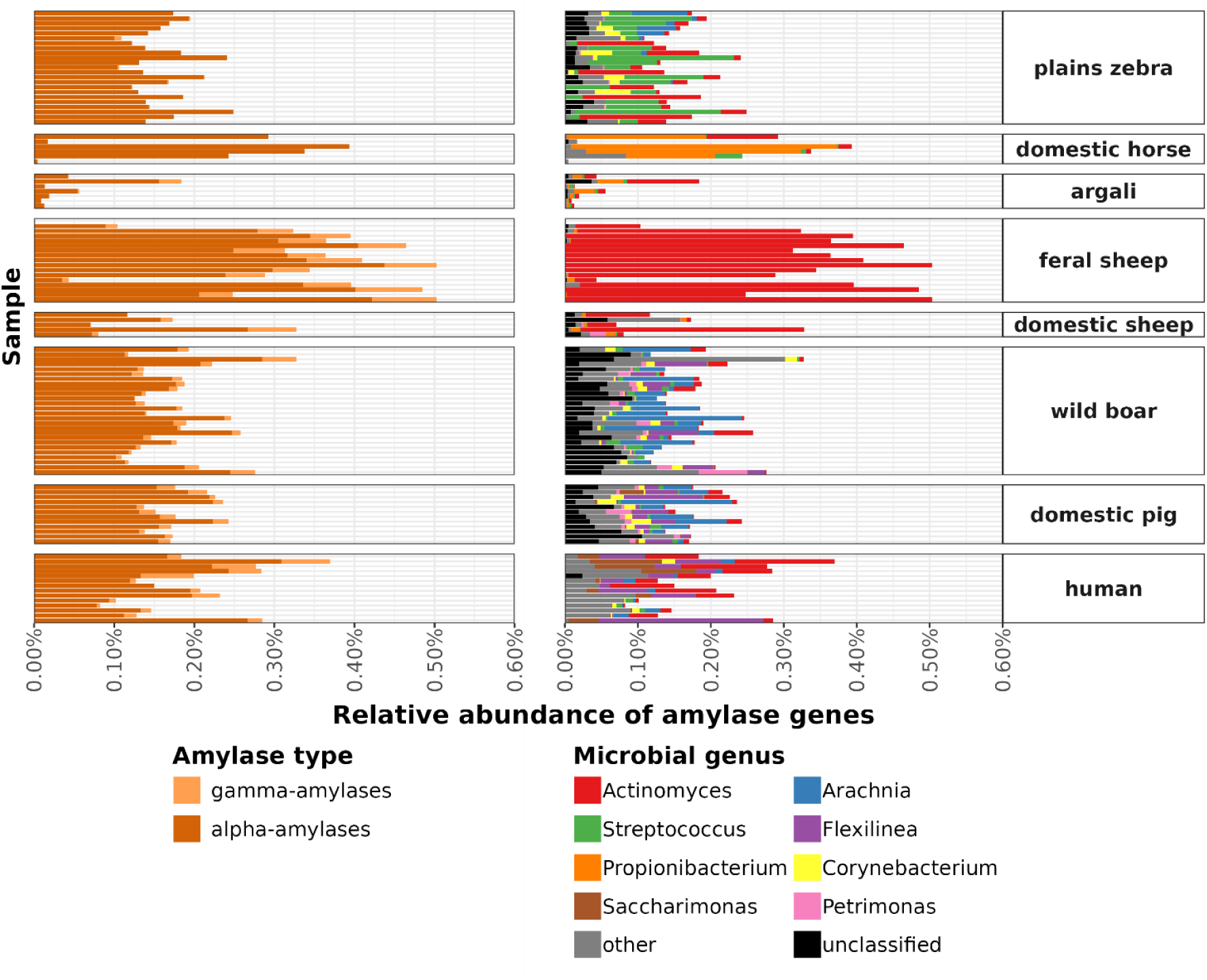
Relative abundances of amylase genes across host species. Left: Bars are coloured by the amylase type. **Right:** Bars are coloured by the bacterial genus encoding the gene. The eight microbial genera accounting for most amylase abundance are shown in distinct colours, while the remaining are labelled as ‘other’. The label ‘unclassified’ indicates genes found on contigs that could not be taxonomically classified at the genus level by CAT.

**Table 3.** Effect of domestication status on the abundance of amylase genes. Tukey HSD results testing for differences in total relative abundance of amylase genes between domestication status categories. This analysis included samples from all genera except humans, with the ‘feral’ category specifically referring to the feral St Kilda sheep. The feral sheep are included here, despite being excluded from other analysis, because they show a markedly increased proportion of amylase genes (See Figure 5).

| Comparison | Difference | Adjusted p-value |
| --- | --- | --- |
| feral-wild | 0.173 | <0.001 |
| domesticated-wild | 0.032 | 0.341 |
| domesticated-feral | -0.141 | <0.001 |

We also investigated the presence of amylase-binding protein abpA, a cell-wall-associated protein encoded by certain oral species of *Streptococcus,* allowing them to utilise host salivary amylases (Nikitkova et al., 2013). Fellows Yates et al. (2021) found increased abundances of abpA genes in the oral microbiome of modern humans compared to other primates, likely as a response to higher starch consumption. As this gene was not present among gene annotations, we identified it based on sequence similarity with publicly available abpA proteins. Beyond reproducing the finding of *Streptococcus*-encoded abpA in modern humans (based on three annotations with high similarity: E-values < 10^-7^), we only found low-confidence hits to abpA in *Desulfobulbus* from domestic sheep (one annotation with E-value = 0.048) and in *Neisseria* from a single pig sample (E-value = 0.011) (Supplementary Figure 9). Owing to the relatively high E-values, the function of these genes cannot be established, especially for the *Desulfobulbus*-encoded sequence, as it is considerably shorter than a typical abpA gene (139 amino acids as opposed to approximately 200). Its similarity to an outer membrane protein suggests the presence of shared domains between the identified gene and abpA (BLAST against RefSeq: E-value = 2e-71, percent identity = 78.4%). Based on these results, the presence of the abpA genes in nonhuman oral microbiomes cannot be established.

### Ancestral microbial lineages are retained in domesticates rather than replaced by human-derived taxa

To understand if domesticates retain their ancestral microorganisms, or if they are colonised by human-derived lineages, we analysed a dataset of 187 MAGs with completeness >75% and contamination <5% (177 bacteria and 10 archaea; Supplementary Tables 15, 16, Supplementary Figures 10, 11). Both a topology– and a distance-based approach were employed. For the topology-based approach we identified nodes that separated, on the one side, a clade containing MAGs from a single domestic-wild pair and, on the other side, a clade containing only human MAGs: we refer to this as a node with a ((domestic, wild), human) topology. We similarly identified ((domestic, human), wild) and ((wild, human), domestic) nodes, which would reflect host-switching events from humans to domestic animals or wild animals, respectively. Finally, to provide a frame of reference, we grouped all nodes that split a clade of MAGs from a single host species and a clade from any other two host species under the label ‘other’. Of the 29 nodes that fit any of the above criteria, seven (24% of 29) showed a ((D, W), H) topology, more frequently than expected by chance (Figure 6A, Supplementary Table 17). Six of the seven ((D, W), H) clades contained MAGs found in the genus *Sus* (Supplementary Figure 12). The one remaining clade was from *Equus*, highlighting that, despite being more phylogenetically and geographically distant than boars and pigs, zebras and domestic horses still carry some highly similar oral symbionts (Figure 6A). No nodes indicating host switching were detected.

**Figure 6.**
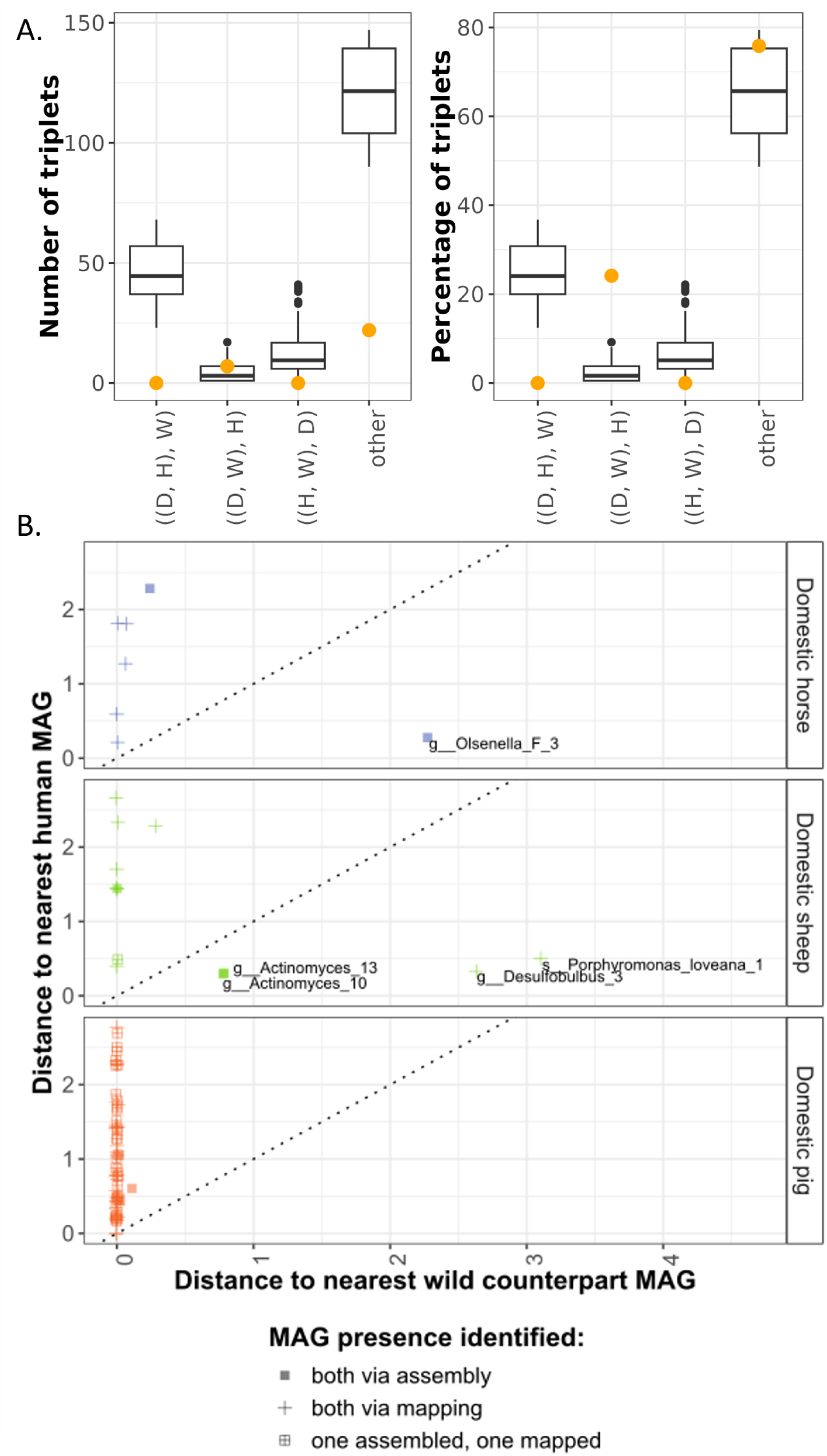
A) Observed (orange point) and permuted (boxplots) counts and percentages of different types of MAG topologies: ((D, H), W) for ((domestic, human), wild); ((D, W), H) for ((domestic, wild), human); ((H, W), D) for ((human, wild), domestic). Note that we required domestic and wild hosts to belong to the same genus. The “other” category includes all other nodes that split MAGs from any single host species on one side, and MAGs from exactly two other host species on the other side. **B)** Comparison of phylogenetic distances between MAG found in domestic animals to the nearest MAG in the respective wild counterpart and to the nearest MAG found in humans. Squares indicate distances that were obtained from the phylogenetic tree of the assembled MAGs, whereas crosses indicate cases where one of the MAGs in the comparison was found through mapping. The dotted parity line represents equal phylogenetic distance to wild and human MAGs. Labelled MAGs show closer phylogenetic distance to a human MAG than to a wild MAG.

The phylogenetic distance-based analysis roughly agreed with the topologies, but in addition detected some putative human-derived microorganisms in sheep and horses. This analysis included 87 MAGs that were found in domestic animals and were in the same genus as another MAG found either in humans or the wild counterpart. Three of these domestic MAGs were identified as present in two different domestic hosts, so they were analysed twice, leading to 90 comparisons in total. Underscoring the extensive similarities within the genus *Sus*, all 70 analysed MAGs from domestic pigs were either also present in boars, or had their closest relative assembled from boars (Figure 6B, Supplementary Table 18). In contrast, though, in sheep we detected genomes of *Actinomyces*, *Desulfobulbus* and *Porphyromonas* (four out of 13 analysed MAGs) that were more closely related to human strains than to any argali strains. However, in none of the cases were the human and domestic genomes each other’s closest relative, with genomes from different hosts found between them in the phylogeny (Supplementary Figure 10), making it unlikely that these indeed represent host-switching events. In horses we identified a more plausible host-switching candidate: one of seven MAGs tested, g Olsenella_F_3, was a sister lineage to s Olsenella_uli_1, a typical oral taxon assembled from humans (Supplementary Figure 10).

In summary, both analyses suggest that ancestral oral microorganisms tend to be retained in domestic animals, and that replacement by human-derived strains is not common. One exception may be *Olsenella*, a typical oral taxon, which not only shows high similarity between domestic horses and humans, but also consistently appears enriched in domestic animals (Figure 2). However, it is important to note that only abundant genomes can be assembled into MAGs, and consequently low-abundance human-derived genomes would have been missed in these analyses.

## Discussion

Several studies exist on the effects of domestication and captivity on animal gut microbiomes (McKenzie et al., 2017; Reese et al., 2021, among others), yet very few have considered other host-associated microbial communities (Podar et al., 2024). Here we investigate differences and similarities in museum-preserved oral microbiomes of three wild-domestic mammalian pairs that represent distinct genera and differ in their diet and time of domestication. Consistent with the domestic (Reese et al., 2021) and captive gut microbiome (McKenzie et al., 2017), host taxonomy had the strongest effect on the taxonomic and functional composition of the oral microbiome, with domestication status showing a smaller but persistent effect (Figure 1C, Supplementary Figure 5A, B) split almost equally between an independent effect, estimated across all host genera, and a host-genus-dependent (interaction) effect (Supplementary Table 7).

Although the three host pairs considered here differ in terms of the evolutionary, geographical and ecological distances between their domestic and wild counterparts, and in the time since domestication (see below), we recover consistent changes that set domestic species apart. Specifically, we identify taxa that are consistently lost, gained, depleted or enriched in all domesticates (Figure 3). Taxa consistently enriched in domesticates, such as *Olsenella* and *Methanobrevibacter*, are also enriched in humans (Figure 3, Supplementary Figure 6). These taxa may drive the observed humanisation of the domesticate oral microbiome (Figure 2), a phenomenon also reported by previous studies of the primate gut microbiome (Clayton et al., 2016; Nishida & Ochman, 2021). Both *Olsenella* and *Methanobrevibacter* are typical members of the human oral microbiome (Dewhirst et al., 2001; Pilliol et al., 2024), with *Methanobrevibacter* especially predominant in the dental calculus of pre-Industrial humans (Gancz et al., 2023; Granehäll et al., 2021). *Methanobrevibacter* is also common in the gut (Samuel et al., 2007), where it has already been associated with captivity in rhinoceroses (Gibson et al., 2019). If confirmed across a wider range of host species, *Olsenella* and *Methanobrevibacter* could serve as ‘domestication indicators’ for the oral microbiome, similar to how *Prevotella* and *Bacterioides* are characteristic of the gut microbiome in captivity (Clayton et al., 2016).

It is important to note that the increased abundance of these taxa in domestic animals does not necessarily reflect colonisation by human-derived microbiota and may instead be driven by shared selective pressures in domesticates and humans, e.g., due to similar diet. In the case of *Olsenella* we found some limited evidence for horizontal transfer from humans: the two most closely related genomes of this taxon were assembled from domestic horses and humans (Supplementary Table 15, Supplementary Figure 10). However, as no *Olsenella* genomes were assembled from zebras, we cannot confirm if the ancestral lineage has been replaced with a human-derived one in horses or if the taxon is naturally absent in zebras (Supplementary Figure 10). For *Methanobrevibacter* only a single MAG was assembled, not allowing the investigation of evolutionary relationships (Supplementary Figure 11). Overall, except for *Olsenella*, our findings show that domestic species retain their microbiota, in agreement with findings from the dog and wolf oral microbiome (Podar et al., 2024), but contrasting with those from the primate gut (Nishida & Ochman, 2021). This suggests that replacement of ancestral microbiota by human strains may be more common in primates than other clades, possibly facilitated by evolutionary relatedness. Finally, it is important to note that, while the MAG-based analysis performed here considers more information than single-gene approaches used by previous studies (Nishida & Ochman, 2021; Podar et al., 2024), it is also inevitably biased towards the most dominant taxa, as those are more likely to be assembled into MAGs and may miss human-derived taxa at low abundances.

Owing to the limitations imposed by what is available in natural history collections, we were unable to obtain samples from direct wild ancestors of domestic sheep and horses, namely the Asian mouflon (*O. gmelini*; Z.-H. Chen et al., 2021; Her et al., 2022) and the Przewalski’s horse (*E. przewalskii*; Gaunitz et al., 2018). This was only possible for the domestic pig, through the inclusion of the wild boar (or Eurasian wild pig). As a result, the phylogenetic distance of domestic-wild pairs differed among the three included genera. The pig was domesticated ∼9K years ago from the wild boar (Giuffra et al., 2000) and the two are considered subspecies by some researchers because they hybridise extensively (Iacolina et al., 2018). If hybridisation leads to exchange of microbiota, it may explain the similarity in microbiome composition between boars and pigs (Figure 1C) and the large number of shared MAGs (Figure 6, Supplementary Figure 12). On the other hand, the domestic sheep and the argali diverged ∼3.5M years ago (Lv et al., 2024), while the domestic horse and zebras diverged ∼4M years ago (Orlando et al., 2009; Vilstrup et al., 2013). While this is a potential caveat, a previous study on the gut microbiome that included both ‘perfect’ wild-domesticated pairs and pairs with more distant wild relatives showed consistent results independent of the phylogenetic distance (Reese et al., 2021). The same study proposed that the effect of domestication may be less pronounced for hosts whose ecology changed less as a result of domestication.

Our results show that the pig microbiome was least affected by domestication (Figures 1C, 2). This could be because they have a more direct wild ancestor included in the comparison, which also shares a geographical range (most domestic pigs and wild pigs included here were from Europe). It is also possible that pigs experiencing smaller change in ecology during domestication compared to other domestic animals, because wild pigs already had a generalist diet. In Central and Eastern Europe, where most of our pig samples come from, it was historically common practice to lead pigs into the forest to find fodder seasonally or year-round (Szabó, 2013; Tóth, 2025). This meant domestic pigs likely consumed a similar diet to wild boars. This of course differs drastically from modern practices, which may explain why the gut microbiome of modern domestic pigs from the USA differed significantly from wild boars, both in terms of taxonomy and functional capacity, reflecting a less diverse diet (Kuthyar et al., 2023).

Reese et al. (2021) also speculated that the effects of domestication may be weaker for more recently domesticated hosts. Our study does not support this idea, as pigs were likely domesticated during the Neolithic period (Vigne et al., 2009), roughly at the same time as sheep (Zeder, 2008), yet we observed stronger differences in sheep than in pigs. However, all factors discussed above, such as phylogenetic divergence, may obscure the relationship between domestication time and microbiome differentiation.

We observe contrasting patterns of microbial richness between wild-domestic pairs (Figure 1A), which is also manifested in abundance shifts and losses of taxa (Figure 3). Specifically, domestic horses showed lower microbiome richness compared to their wild counterparts, whereas domestic pigs and sheep tend towards higher richness (albeit not significant; Figure 1A, Table 2, Supplementary Table 5). A potential caveat may be the very different geographical origins for horses and zebras: horse specimens were all of European breeds, whereas plains zebras were from sub-Saharan Africa (Supplementary Table 1). However, previous research reported that geographical location had a much smaller impact on the microbiome than domestication or captivity status (Clayton et al., 2016), while specifically latitude showed no impact on oral microbiome richness (Moraitou et al., 2026). In addition, our results on microbiome richness agree with previous work reporting higher gut microbiome richness in Przewalski’s horses than domestic horses (Metcalf et al., 2017), but comparable richness between domestic pigs with wild boars (Kuthyar et al., 2023). We also highlight that, as has already been shown for the gut microbiome (McKenzie et al., 2017; Reese et al., 2021), domestication does not always result in less diverse oral microbiota and that the effects may differ from species to species.

Along with taxonomic differences, we also detected consistent differences in the functional potential of the oral microbiome in domesticates compared to wild species (Figure 4A, Figure 5, Supplementary Figure 7). Specifically, several pathways showed lower abundance in all three domestic animals (Figure 4B). This deviates from a comparison of the gut microbiomes of domestic pigs and wild boars, which found the domesticates to have a larger functional diversity (Kuthyar et al., 2023). In horses and sheep, we found a significant depletion in the starch and sucrose metabolism pathway, and especially a lower abundance of the beta-glucosidase gene, which catalyses the degradation of cellulose. In combination with the trend towards increased amylase abundance in these host species (Figure 5, Table 3), this observation suggests that the oral microbiome may have adapted to a reduced fibre and increased starch intake in domesticated animals. Once again, wild boars and pigs showed fewer differences in functional capacity (Figures 4 and 5), which is consistent with a less pronounced change in diet with domestication.

Finally, the feral sheep of St Kilda provided an opportunity to investigate whether, and to what extent, the impact of domestication is reversible. Reverting to a wild lifestyle seemed to partially restore a wild-like oral microbiome. The feral sheep oral microbiome was more similar to that of the argali than to that of the domestic sheep, both in terms of richness and taxonomic composition (Figures 1, 2). Even the abundances of the domestic-associated *Olsenella* and *Methanobrevibacter* and the wild-associated *Mannheimia* and *Basfia* were similar in feral sheep and argalis, whereas abundances of other taxa were intermediate between wild and domestic (Supplementary Figure 6). Surprisingly, though, the feral sheep of St Kilda showed a marked increase in microbial amylase genes, compared to both fully domestic and wild counterparts (Figure 5, Table 3). These sheep rely on local grasses and shrubs for their diet, so it is unclear what drove this increase in amylase genes, but it may be a remnant of their domesticated past.

In conclusion, we find that the oral microbiome of domestic animals differs from that of their wild counterparts both in terms of taxonomic composition and functional capacity, generally reflecting increased proximity to humans and adaptation to increased starch consumption. These differences are shared by all three host genera, despite inherent differences in their ecology, digestion and dietary class. However, the degree of change differs among the host pairs considered here, reflecting a multitude of factors, like the extent of dietary differences and the time since phylogenetic divergence between domestic and wild counterparts, as proposed for the pig-boar pair. Despite the increased abundance of human-associated taxa in domestic mammals of all three pairs (including pigs), we do not find extensive replacement of ancestral wild microorganisms by human lineages.

## Acknowledgements

We would like to express our gratitude to Germán Hernández Alonso (University of Edinburgh) for his contribution to bioinformatic analysis and to Phaedra Kokkini, Richard Sabin, Roberto Portela Miguez (Natural History Museum London) and Frank Zachos (Natural History Museum Vienna) for their support during sampling. This work was supported by the Swedish Research Council (Formas) grant number 2019-00275 to KG.

## Data Availability Note

The scripts used for this analysis have been deposited at https://github.com/markella-moraitou/domesticated_oral_mb.

## References

1. Aarestrup, F. M., Butaye, P., & Witte, W. (2002). Nonhuman Reservoirs of Enterococci. In The Enterococci (pp. 55–99). John Wiley & Sons, Ltd. 10.1128/9781555817923.ch2

2. Abranches, J., Zeng, L., Kajfasz, J. K., Palmer, S. R., Chakraborty, B., Wen, Z. T., Richards, V. P., Brady, L. J., & Lemos, J. A. (2018). Biology of Oral Streptococci. Microbiology Spectrum, 6(5), 10.1128/microbiolspec.gpp3-0042-2018. https://doi.org/10.1128/microbiolspec.gpp3-0042-2018

3. Aitchison, J. W. (1986). The Statistical Analysis of Compositional Data. Springer Netherlands.

4. Alberdi, A. (2025). distillR: R package for distilling functional annotations of bacterial genomes and metagenomes (Version 0.3.0) [Computer software]. https://github.com/anttonalberdi/distillR

5. Alberdi, A., Aizpurua, O., Bohmann, K., Zepeda-Mendoza, M. L., & Gilbert, M. T. P. (2016). Do Vertebrate Gut Metagenomes Confer Rapid Ecological Adaptation? Trends in Ecology & Evolution, 31(9), 689–699. 10.1016/j.tree.2016.06.008

6. Angen, Ø., Mutters, R., Caugant, D. A., Olsen, J. E., & Bisgaard, M. (1999). Taxonomic relationships of the [Pasteurella] haemolytica complex as evaluated by DNA-DNA hybridizations and 16S rRNA sequencing with proposal of Mannheimia haemolytica gen. Nov., comb. Nov., Mannheimia granulomatis comb. Nov., Mannheimia glucosida sp. Nov., Mannheimia ruminalis sp. Nov. And Mannheimia varigena sp. Nov. International Journal of Systematic and Evolutionary Microbiology, 49(1), 67–86. 10.1099/00207713-49-1-67

7. Asangba, A. E., Mugisha, L., Rukundo, J., Lewis, R. J., Halajian, A., Cortés-Ortiz, L., Junge, R. E., Irwin, M. T., Karlson, J., Perkin, A., Watsa, M., Erkenswick, G., Bales, K. L., Patton, D. L., Jasinska, A. J., Fernandez-Duque, E., Leigh, S. R., & Stumpf, R. M. (2022). Large Comparative Analyses of Primate Body Site Microbiomes Indicate that the Oral Microbiome Is Unique among All Body Sites and Conserved among Nonhuman Primates. Microbiology Spectrum, 10(3), e01643–21. 10.1128/spectrum.01643-21

8. Bachmann, B. J. (1955). Studies on Cytophaga fermentans, n.sp., a Facultatively Anaerobic Lower Myxobacterium. Microbiology, 13(3), 541–551. 10.1099/00221287-13-3-541

9. Bae, D.-Y., Moon, S.-H., Lee, T. G., Ko, Y.-S., Cho, Y.-C., Kang, H., Park, C.-S., Kang, J.-S., Oh, Y., & Cho, H.-S. (2025). Consequences of Domestication on Gut Microbiome: A Comparative Analysis Between Wild Boars and Domestic Pigs. Animals, 15(5). 10.3390/ani15050747

10. Barnett, D. J. m, Arts, I. C. w, & Penders, J. (2021). microViz: An R package for microbiome data visualization and statistics. Journal of Open Source Software, 6(63), 3201. 10.21105/joss.03201

11. Bowers, R. M., The Genome Standards Consortium, Kyrpides, N. C., Stepanauskas, R., Harmon-Smith, M., Doud, D., Reddy, T. B. K. K., Schulz, F., Jarett, J., Rivers, A. R., Eloe-Fadrosh, E. A., Tringe, S. G., Ivanova, N. N., Copeland, A., Clum, A., Becraft, E. D., Malmstrom, R. R., Birren, B., Podar, M., … Woyke, T. (2017). Minimum information about a single amplified genome (MISAG) and a metagenome-assembled genome (MIMAG) of bacteria and archaea. Nature Biotechnology, 35(8), 725–731. 10.1038/nbt.3893

12. Boyd, J. M., Doney, J. M., Gunn, R. G., & Jewell, P. A. (1964). The Soay Sheep of the Island of Hirta, St. Kilda. A Study of a Feral Population. Proceedings of the Zoological Society of London, 142(1), 129–164. 10.1111/j.1469-7998.1964.tb05159.x

13. Bushnell, B. (2014). BBMap: A fast, accurate, splice-aware aligner (Issue LBNL-7065E). Lawrence Berkeley National Lab. (LBNL), Berkeley, CA (United States).

14. Chaumeil, P.-A., Mussig, A. J., Hugenholtz, P., & Parks, D. H. (2022). GTDB-Tk v2: Memory friendly classification with the genome taxonomy database. Bioinformatics, 38(23), 5315–5316. 10.1093/bioinformatics/btac672

15. Chen, S., Zhou, Y., Chen, Y., & Gu, J. (2018). fastp: An ultra-fast all-in-one FASTQ preprocessor. Bioinformatics, 34(17), i884–i890.

16. Chen, Z.-H., Xu, Y.-X., Xie, X.-L., Wang, D.-F., Aguilar-Gómez, D., Liu, G.-J., Li, X., Esmailizadeh, A., Rezaei, V., Kantanen, J., Ammosov, I., Nosrati, M., Periasamy, K., Coltman, D. W., Lenstra, J. A., Nielsen, R., & Li, M.-H. (2021). Whole-genome sequence analysis unveils different origins of European and Asiatic mouflon and domestication-related genes in sheep. Communications Biology, 4(1), 1307. 10.1038/s42003-021-02817-4

17. Clausen, P. T. L. C., Aarestrup, F. M., & Lund, O. (2018). Rapid and precise alignment of raw reads against redundant databases with KMA. BMC Bioinformatics, 19(1), 307. 10.1186/s12859-018-2336-6

18. Clayton, J. B., Vangay, P., Huang, H., Ward, T., Hillmann, B. M., Al-Ghalith, G. A., Travis, D. A., Long, H. T., Van Tuan, B., VanMinh, V., Cabana, F., Nadler, T., Toddes Barbaraand Murphy, T., Glander, K. E., & Johnson Timothy Jand Knights, D. (2016). Captivity humanizes the primate microbiome. Proc. Natl. Acad. Sci. U. S. A., 113(37), 10376–10381.

19. Dallas, J. W., & Warne, R. W. (2023). Captivity and Animal Microbiomes: Potential Roles of Microbiota for Influencing Animal Conservation. Microbial Ecology, 85(3), 820–838. 10.1007/s00248-022-01991-0

20. Danecek, P., Bonfield, J. K., Liddle, J., Marshall, J., Ohan, V., Pollard, M. O., Whitwham, A., Keane, T., McCarthy, S. A., Davies, R. M., & Li, H. (2021). Twelve years of SAMtools and BCFtools. Gigascience, 10(2).

21. Darwin, C. (1868). The Variation of Animals and Plants Under Domestication. J. Murray.

22. Dewhirst, F. E., Paster, B. J., Tzellas, N., Coleman, B., Downes, J., Spratt, D. A., & Wade, W. G. (2001). Characterization of novel human oral isolates and cloned 16S rDNA sequences that fall in the family Coriobacteriaceae: Description of olsenella gen. Nov., reclassification of Lactobacillus uli as Olsenella uli comb. Nov. And description of Olsenella profusa sp. Nov. Int. J. Syst. Evol. Microbiol., 51(Pt 5), 1797–1804.

23. Duitama González, C., Vicedomini, R., Lemane, T., Rascovan, N., Richard, H., & Chikhi, R. (2023). decOM: Similarity-based microbial source tracking of ancient oral samples using k-mer-based methods. Microbiome, 11(1), 243. 10.1186/s40168-023-01670-3

24. Eddy, S. R. (2009). A new generation of homology search tools based on probabilistic inference. In Genome Informatics 2009 (pp. 205–211). PUBLISHED BY IMPERIAL COLLEGE PRESS AND DISTRIBUTED BY WORLD SCIENTIFIC PUBLISHING CO. 10.1142/9781848165632_0019

25. Farese, M., Moraitou, M., Jin, C., Forsythe, A., Micarelli, I., Valk, T. van der, Manzi, G., Parducci, L., Tafuri, M. A., & Guschanski, K. (2026). Lifestyle Impacts the Oral Microbiome of Classical and Post-Classical Societies in Italy. American Journal of Biological Anthropology, 191(1), e70357. 10.1002/ajpa.70357

26. Fatoretto, B. T., Gonzalez, I. H. L., Lima, C. F. d. M., Monticelli, C., & Ramos, P. L. (2022). A comparison of rectal and oral cultivable microbiota in wild and captive black lion tamarins (Leontopithecus chrysopygus, Mikan 1823). American Journal of Primatology, 84(4–5), e23370. 10.1002/ajp.23370

27. Fellows Yates, J. A., Velsko, I. M., Aron, F., Posth, C., Hofman, C. A., Austin, R. M., Parker, C. E., Mann, A. E., Nägele, K., Arthur, K. W., Arthur, J. W., Bauer, C. C., Crevecoeur, I., Cupillard, C., Curtis, M. C., Dalén, L., Díaz-Zorita Bonilla, M., Díez Fernández-Lomana, J. C., Drucker, D. G., … Warinner, C. (2021). The evolution and changing ecology of the African hominid oral microbiome. Proceedings of the National Academy of Sciences, 118(20), e2021655118. 10.1073/pnas.2021655118

28. Gancz, A. S., Farrer, A. G., Nixon, M. P., Wright, S., Arriola, L., Adler, C., Davenport, E. R., Gully, N., Cooper, A., Britton, K., Dobney, K., Silverman, J. D., & Weyrich, L. S. (2023). Ancient dental calculus reveals oral microbiome shifts associated with lifestyle and disease in Great Britain. Nature Microbiology, 8(12), 2315–2325. 10.1038/s41564-023-01527-3

29. Gaunitz, C., Fages, A., Hanghøj, K., Albrechtsen, A., Khan, N., Schubert, M., Seguin-Orlando, A., Owens, I. J., Felkel, S., Bignon-Lau, O., de Barros Damgaard, P., Mittnik, A., Mohaseb, A. F., Davoudi, H., Alquraishi, S., Alfarhan, A. H., Al-Rasheid, K. A. S., Crubézy, E., Benecke, N., … Orlando, L. (2018). Ancient genomes revisit the ancestry of domestic and Przewalski’s horses. Science, 360(6384), 111–114. 10.1126/science.aao3297

30. Gearty, W., & Jones, L. A. (2023). rphylopic: An R package for fetching, transforming, and visualising PhyloPic silhouettes. Methods in Ecology and Evolution, 14(11), 2700–2708. 10.1111/2041-210X.14221

31. Gibson, K. M., Nguyen, B. N., Neumann, L. M., Miller, M., Buss, P., Daniels, S., Ahn, M. J., Crandall, K. A., & Pukazhenthi, B. (2019). Gut microbiome differences between wild and captive black rhinoceros – implications for rhino health. Scientific Reports, 9(1), 7570. 10.1038/s41598-019-43875-3

32. Gilroy, R., Leng, J., Ravi, A., Adriaenssens, E. M., Oren, A., Baker, D., Ragione, R. M. L., Proudman, C., & Pallen, M. J. (2022). Metagenomic investigation of the equine faecal microbiome reveals extensive taxonomic diversity. PeerJ, 10, e13084. 10.7717/peerj.13084

33. Giuffra, E., Kijas, J. M. H., Amarger, V., Carlborg, Ö., Jeon, J.-T., & Andersson, L. (2000). The Origin of the Domestic Pig: Independent Domestication and Subsequent Introgression. Genetics, 154(4), 1785–1791. 10.1093/genetics/154.4.1785

34. Gloor, G. B., Macklaim, J. M., Pawlowsky-Glahn, V., & Egozcue, J. J. (2017). Microbiome Datasets Are Compositional: And This Is Not Optional. 8, 2224. 10.3389/fmicb.2017.02224

35. Granehäll, L., Huang, K. D., Tett, A., Manghi, P., Paladin, A., O’Sullivan, N., Rota-Stabelli, O., Segata, N., Zink, A., & Maixner, F. (2021). Metagenomic analysis of ancient dental calculus reveals unexplored diversity of oral archaeal Methanobrevibacter. Microbiome 2021 9:1, 9(1), 1–18. 10.1186/S40168-021-01132-8

36. Hadfield, J. D. (2010). MCMC Methods for Multi-Response Generalized Linear Mixed Models: The MCMCglmm R Package. Journal of Statistical Software, 33, 1–22. 10.18637/jss.v033.i02

37. Hammer, K. (1984). Das Domestikationssyndrom. Die Kulturpflanze, 32(1), 11–34. 10.1007/BF02098682

38. Hauser, M., Steinegger, M., & Söding, J. (2016). MMseqs software suite for fast and deep clustering and searching of large protein sequence sets. Bioinformatics, 32(9), 1323–1330. 10.1093/bioinformatics/btw006

39. Hedlund, B. P., & Tønjum, T. (2015). Simonsiella. In Bergey’s Manual of Systematics of Archaea and Bacteria (pp. 1–12). John Wiley & Sons, Ltd. 10.1002/9781118960608.gbm00983

40. Her, C., Rezaei, H.-R., Hughes, S., Naderi, S., Duffraisse, M., Mashkour, M., Naghash, H.-R., Bălășescu, A., Luikart, G., Jordan, S., Özüt, D., Kence, A., Bruford, M. W., Tresset, A., Vigne, J.-D., Taberlet, P., Hänni, C., & Pompanon, F. (2022). Broad maternal geographic origin of domestic sheep in Anatolia and the Zagros. Animal Genetics, 53(3), 452–459. 10.1111/age.13191

41. Iacolina, L., Pertoldi, C., Amills, M., Kusza, S., Megens, H.-J., Bâlteanu, V. A., Bakan, J., Cubric-Curik, V., Oja, R., Saarma, U., Scandura, M., Šprem, N., & Stronen, A. V. (2018). Hotspots of recent hybridization between pigs and wild boars in Europe. Scientific Reports, 8(1), 17372. 10.1038/s41598-018-35865-8

42. Julliand, V., De Fombelle, A., & Varloud, M. (2006). Starch digestion in horses: The impact of feed processing. *Livestock Science*, Nutritive Value of Concentrates in Horses, 100(1), 44–52. 10.1016/j.livprodsci.2005.11.001

43. Juni, E., & Heym, G. A. (1986). Psychrobacter immobilis gen. nov., sp. nov.: Genospecies Composed of Gram-Negative, Aerobic, Oxidase-Positive Coccobacilli. International Journal of Systematic and Evolutionary Microbiology, 36(3), 388–391. 10.1099/00207713-36-3-388

44. Krakau, S., Straub, D., Gourlé, H., Gabernet, G., & Nahnsen, S. (2022). nf-core/mag: A best-practice pipeline for metagenome hybrid assembly and binning. NAR Genomics and Bioinformatics, 4(1), lqac007. 10.1093/nargab/lqac007

45. Kuhnert, P., Scholten, E., Haefner, S., Mayor, D., & Frey, J. (2010). Basfia succiniciproducens gen. Nov., sp. Nov., a new member of the family Pasteurellaceae isolated from bovine rumen. International Journal of Systematic and Evolutionary Microbiology, 60(Pt 1), 44–50. 10.1099/ijs.0.011809-0

46. Kuthyar, S., Diaz, J., Avalos-Villatoro, F., Maltecca, C., Tiezzi, F., Dunn, R. R., & Reese, A. T. (2023). Domestication shapes the pig gut microbiome and immune traits from the scale of lineage to population. Journal of Evolutionary Biology, 36(12), 1695–1711. 10.1111/jeb.14227

47. Li, D., Luo, R., Liu, C. M., Leung, C. M., Ting, H. F., Sadakane, K., Yamashita, H., & Lam, T. W. (2016). MEGAHIT v1.0: A fast and scalable metagenome assembler driven by advanced methodologies and community practices. Methods, 102, 3–11. 10.1016/j.ymeth.2016.02.020

48. Li, H., & Durbin, R. (2009). Fast and accurate short read alignment with Burrows-Wheeler transform. Bioinformatics, 25(14), 1754–1760.

49. Librado, P., Khan, N., Fages, A., Kusliy, M. A., Suchan, T., Tonasso-Calvière, L., Schiavinato, S., Alioglu, D., Fromentier, A., Perdereau, A., Aury, J.-M., Gaunitz, C., Chauvey, L., Seguin-Orlando, A., Der Sarkissian, C., Southon, J., Shapiro, B., Tishkin, A. A., Kovalev, A. A., … Orlando, L. (2021). The origins and spread of domestic horses from the Western Eurasian steppes. Nature, 598(7882), 634–640. 10.1038/s41586-021-04018-9

50. Lin, H., & Peddada, S. D. (2024). Multigroup analysis of compositions of microbiomes with covariate adjustments and repeated measures. Nature Methods, 21(1), 83–91. 10.1038/s41592-023-02092-7

51. Lunesu, M. F., Decandia, M., Molle, G., Atzori, A. S., Bomboi, G. C., & Cannas, A. (2021). Dietary Starch Concentration Affects Dairy Sheep and Goat Performances Differently during Mid-Lactation. Animals, 11(5). 10.3390/ani11051222

52. Lv, F.-H., Wang, D.-F., Zhao, S.-Y., Lv, X.-Y., Sun, W., Nielsen, R., & Li, M.-H. (2024). Deep Ancestral Introgressions between Ovine Species Shape Sheep Genomes via Argali-Mediated Gene Flow. Molecular Biology and Evolution, 41(11), msae212. 10.1093/molbev/msae212

53. Madden, T. (2003). The BLAST Sequence Analysis Tool. In The NCBI Handbook [Internet]. National Center for Biotechnology Information (US). https://www.ncbi.nlm.nih.gov/books/NBK21097/

54. McKenzie, V. J., Song, S. J., Delsuc, F., Prest, T. L., Oliverio, A. M., Korpita, T. M., Alexiev, A., Amato, K. R., Metcalf, J. L., Kowalewski, M., Avenant, N. L., Link, A., Di Fiore, A., Seguin-Orlando, A., Feh, C., Orlando, L., Mendelson, J. R., Sanders, J., & Knight, R. (2017). The Effects of Captivity on the Mammalian Gut Microbiome. Integrative and Comparative Biology, 57(4), 690–704. 10.1093/icb/icx090

55. McMurdie, P. J., & Holmes, S. (2013). phyloseq: An R Package for Reproducible Interactive Analysis and Graphics of Microbiome Census Data. PLoS ONE, 8(4), e61217.

56. Metcalf, J. L., Song, S. J., Morton, J. T., Weiss, S., Seguin-Orlando, A., Joly, F., Feh, C., Taberlet, P., Coissac, E., Amir, A., Willerslev, E., Knight, R., McKenzie, V., & Orlando, L. (2017). Evaluating the impact of domestication and captivity on the horse gut microbiome. Scientific Reports, 7(1), Article 1. 10.1038/s41598-017-15375-9

57. Moraitou, M., Richards, J. L., Bolyos, C., Saliari, K., Gilissen, E., Timmons, Z., Kitchener, A. C., Pauwels, O. S. G., Sabin, R., Kokkini, P., Portela Miguez, R., & Guschanski, K. (2025). Host Traits Impact the Outcome of Metagenomic Library Preparation From Dental Calculus Samples Across Diverse Mammals. Molecular Ecology Resources, 25(8), e70039. 10.1111/1755-0998.70039

58. Moraitou, M., Rozwalak, P., Richards, J., Saliari, K., Gilissen, E., Timmons, Z., Kitchener, A. C., Pauwels, O. S. G., Sabin, R., Kokkini, P., Miguez, R. P., Kalthoff, D., Kellner, F. L., Brealey, J. C., Dutilh, B. E., Martin, M. D., Veiberg, V., & Guschanski, K. (2026). Museum specimens reveal evolutionary drivers and functions of oral microbiome in mammals (p. 2026.06.19.733412). bioRxiv. 10.64898/2026.06.19.733412

59. Narat, V., Amato, K. R., Ranger, N., Salmona, M., Mercier-Delarue, S., Rupp, S., Ambata, P., Njouom, R., Simon, F., Giles-Vernick, T., & LeGoff, J. (2020). A multi-disciplinary comparison of great ape gut microbiota in a central African forest and European zoo. Scientific Reports, 10(1), 19107. 10.1038/s41598-020-75847-3

60. Nikitkova, A. E., Haase, E. M., & Scannapieco, F. A. (2013). Taking the Starch out of Oral Biofilm Formation: Molecular Basis and Functional Significance of Salivary α-Amylase Binding to Oral Streptococci. Applied and Environmental Microbiology, 79(2), 416–423. 10.1128/AEM.02581-12

61. Nishida, A. H., & Ochman, H. (2021). Captivity and the co-diversification of great ape microbiomes. Nature Communications, 12(1), 5632. 10.1038/s41467-021-25732-y

62. Oksanen, J., Simpson, G. L., Blanchet, F. G., Kindt, R., Legendre, P., Minchin, P. R., O’Hara, R. B., Solymos, P., Stevens, M. H. H., Szoecs, E., Wagner, H., Barbour, M., Bedward, M., Bolker, B., Borcard, D., Carvalho, G., Chirico, M., De Caceres, M., Durand, S., … Weedon, J. (2022). Vegan: Community Ecology Package. R package version 2.6-2. 2.5-6.

63. Olm, M. R., Brown, C. T., Brooks, B., & Banfield, J. F. (2017). dRep: A tool for fast and accurate genomic comparisons that enables improved genome recovery from metagenomes through de-replication. ISME J., 11(12), 2864–2868.

64. Orlando, L., Metcalf, J. L., Alberdi, M. T., Telles-Antunes, M., Bonjean, D., Otte, M., Martin, F., Eisenmann, V., Mashkour, M., Morello, F., Prado, J. L., Salas-Gismondi, R., Shockey, B. J., Wrinn, P. J., Vasil’ev, S. K., Ovodov, N. D., Cherry, M. I., Hopwood, B., Male, D., … Cooper, A. (2009). Revising the recent evolutionary history of equids using ancient DNA. Proceedings of the National Academy of Sciences, 106(51), 21754–21759. 10.1073/pnas.0903672106

65. Pajic, P., Pavlidis, P., Dean, K., Neznanova, L., Romano, R.-A., Garneau, D., Daugherity, E., Globig, A., Ruhl, S., & Gokcumen, O. (2019). Independent amylase gene copy number bursts correlate with dietary preferences in mammals. eLife, 8, e44628. 10.7554/eLife.44628

66. Parks, D. H., Chuvochina, M., Rinke, C., Mussig, A. J., Chaumeil, P.-A., & Hugenholtz, P. (2022). GTDB: An ongoing census of bacterial and archaeal diversity through a phylogenetically consistent, rank normalized and complete genome-based taxonomy. Nucleic Acids Research, 50(D1), D785–D794. 10.1093/nar/gkab776

67. Parks, D. H., Imelfort, M., Skennerton, C. T., Hugenholtz, P., & Tyson, G. W. (2015). CheckM: Assessing the quality of microbial genomes recovered from isolates, single cells, and metagenomes. Genome Research, 25(7), 1043–1055. 10.1101/gr.186072.114

68. Pilliol, V., Mahmoud Abdelwadoud, B., Aïcha, H., Lucille, T., Gérard, A., Hervé, T., Michel, D., Ghiles, G., & Elodie, T. (2024). Methanobrevibacter oralis: A comprehensive review. Journal of Oral Microbiology, 16(1), 2415734. 10.1080/20002297.2024.2415734

69. Pimiento, C., & Antonelli, A. (2022). Integrating deep-time palaeontology in conservation prioritisation. Frontiers in Ecology and Evolution, 10. https://www.frontiersin.org/articles/10.3389/fevo.2022.959364

70. Podar, N. A., Carrell, A. A., Cassidy, K. A., Klingeman, D. M., Yang, Z., Stahler, E. A., Smith, D. W., Stahler, D. R., & Podar, M. (2024). From wolves to humans: Oral microbiome resistance to transfer across mammalian hosts. mBio, 15(3), e03342–23. 10.1128/mbio.03342-23

71. Prabhu, V. R., Wasimuddin, Kamalakkannan, R., Arjun, M. S., & Nagarajan, M. (2020). Consequences of Domestication on Gut Microbiome: A Comparative Study Between Wild Gaur and Domestic Mithun. Frontiers in Microbiology, 11. https://www.frontiersin.org/articles/10.3389/fmicb.2020.00133

72. Reese, A. T., Chadaideh, K. S., Diggins, C. E., Schell, L. D., Beckel, M., Callahan, P., Ryan, R., Emery Thompson, M., & Carmody, R. N. (2021). Effects of domestication on the gut microbiota parallel those of human industrialization. eLife, 10, e60197. 10.7554/eLife.60197

73. Samuel, B. S., Hansen, E. E., Manchester, J. K., Coutinho, P. M., Henrissat, B., Fulton, R., Latreille, P., Kim, K., Wilson, R. K., & Gordon, J. I. (2007). Genomic and metabolic adaptations of Methanobrevibacter smithii to the human gut. Proceedings of the National Academy of Sciences, 104(25), 10643–10648. 10.1073/pnas.0704189104

74. Sanders, J. G., Sprockett, D. D., Li, Y., Mjungu, D., Lonsdorf, E. V., Ndjango, J.-B. N., Georgiev, A. V., Hart, J. A., Sanz, C. M., Morgan, D. B., Peeters, M., Hahn, B. H., & Moeller, A. H. (2023). Widespread extinctions of co-diversified primate gut bacterial symbionts from humans. Nature Microbiology, 1–12. 10.1038/s41564-023-01388-w

75. Shaffer, M., Borton, M. A., McGivern, B. B., Zayed, A. A., La Rosa, S. L., Solden, L. M., Liu, P., Narrowe, A. B., Rodríguez-Ramos, J., Bolduc, B., Gazitúa, M. C., Daly, R. A., Smith, G. J., Vik, D. R., Pope, P. B., Sullivan, M. B., Roux, S., & Wrighton, K. C. (2020). DRAM for distilling microbial metabolism to automate the curation of microbiome function. Nucleic Acids Research, 48(16), 8883–8900. 10.1093/nar/gkaa621

76. Shaiber, A., Willis, A. D., Delmont, T. O., Roux, S., Chen, L. X., Schmid, A. C., Yousef, M., Watson, A. R., Lolans, K., Esen, Ö. C., Lee, S. T. M., Downey, N., Morrison, H. G., Dewhirst, F. E., Mark Welch, J. L., & Eren, A. M. (2020). Functional and genetic markers of niche partitioning among enigmatic members of the human oral microbiome. Genome Biology, 21(1), 1–35. 10.1186/s13059-020-02195-w

77. Shanmugam, M., Abbar, F. E., & Ramasubbu, N. (2015). Transcriptome Profiling of Wild-Type and pga-Knockout Mutant Strains Reveal the Role of Exopolysaccharide in Aggregatibacter actinomycetemcomitans. PLOS ONE, 10(7), e0134285. 10.1371/journal.pone.0134285

78. Silverman, J. D., Washburne, A. D., Mukherjee, S., & David, L. A. (2017). A phylogenetic transform enhances analysis of compositional microbiota data. eLife, 6, e21887. 10.7554/eLife.21887

79. Suzuki, T. A., & Ley, R. E. (2020). The role of the microbiota in human genetic adaptation. Science, 370(6521), eaaz6827. 10.1126/science.aaz6827

80. Sweeny, A. R., Lemon, H., Ibrahim, A., Watt, K. A., Wilson, K., Childs, D. Z., Nussey, D. H., Free, A., & McNally, L. (2023). A mixed-model approach for estimating drivers of microbiota community composition and differential taxonomic abundance. mSystems, 8(4), e00040–23. 10.1128/msystems.00040-23

81. Szabó, P. (2013). Rethinking pannage: Historical interactions between oak and swine. In Trees, Forested Landscapes and Grazing Animals. Routledge.

82. Tenenbaum, D., & Maintainer, B. P. (2024). KEGGREST: Client-side REST access to the Kyoto Encyclopedia of Genes and Genomes (*KEGG* [Computer software]. https://bioconductor.org/packages/KEGGREST

83. Tóth, K. (2025). Oinking in the Mountains: Pigs and Pannage in the Principality of Transylvania (1541–1699). Animal History, 2(1), 60–79. 10.1525/ah.2025.2709842

84. Velsko, I. M., Fellows Yates, J. A., Aron, F., Hagan, R. W., Frantz, L. A. F., Loe, L., Martinez, J. B. R., Chaves, E., Gosden, C., Larson, G., & Warinner, C. (2019). Microbial differences between dental plaque and historic dental calculus are related to oral biofilm maturation stage. Microbiome, 7(1), 102. 10.1186/s40168-019-0717-3

85. Vigne, J.-D., Zazzo, A., Saliège, J.-F., Poplin, F., Guilaine, J., & Simmons, A. (2009). Pre-Neolithic wild boar management and introduction to Cyprus more than 11,400 years ago. Proceedings of the National Academy of Sciences of the United States of America, 106(38), 16135–16138. 10.1073/pnas.0905015106

86. Vilstrup, J. T., Seguin-Orlando, A., Stiller, M., Ginolhac, A., Raghavan, M., Nielsen, S. C. A., Weinstock, J., Froese, D., Vasiliev, S. K., Ovodov, N. D., Clary, J., Helgen, K. M., Fleischer, R. C., Cooper, A., Shapiro, B., & Orlando, L. (2013). Mitochondrial Phylogenomics of Modern and Ancient Equids. PLOS ONE, 8(2), e55950. 10.1371/journal.pone.0055950

87. von Meijenfeldt, F. A. B., Arkhipova, K., Cambuy, D. D., Coutinho, F. H., & Dutilh, B. E. (2019). Robust taxonomic classification of uncharted microbial sequences and bins with CAT and BAT. Genome Biology, 20(1), 217. 10.1186/s13059-019-1817-x

88. Voolstra, C. R., Hermann, R. J., Stibor, H., Keller, A., & Becks, L. (2026). Timescale mismatch: Redefining ecological opportunity for evolution in the Anthropocene. The EMBO Journal, 45(18), 6359–6366. 10.1038/s44318-026-00880-3

89. Wilkins, A. S., Wrangham, R. W., & Fitch, W. T. (2014). The “Domestication Syndrome” in Mammals: A Unified Explanation Based on Neural Crest Cell Behavior and Genetics. Genetics, 197(3), 795–808. 10.1534/genetics.114.165423

90. Yang, S.-H., Seo, H.-S., Woo, J.-H., Oh, H.-M., Jang, H., Lee, J.-H., Kim, S.-J., & Kwon, K. K. (2014). Carboxylicivirga gen. Nov. In the family Marinilabiliaceae with two novel species, Carboxylicivirga mesophila sp. Nov. And Carboxylicivirga taeanensis sp. Nov., and reclassification of Cytophaga fermentans as Saccharicrinis fermentans gen. Nov., comb. Nov. International Journal of Systematic and Evolutionary Microbiology, 64(Pt_4), 1351–1358. 10.1099/ijs.0.053462-0

91. Zeder, M. A. (2008). Domestication and early agriculture in the Mediterranean Basin: Origins, diffusion, and impact. Proceedings of the National Academy of Sciences, 105(33), 11597–11604. 10.1073/pnas.0801317105

