## Supplementary Figures for "Shared and genus-specific impact of domestication on mammal oral microbiomes revealed through museomics"

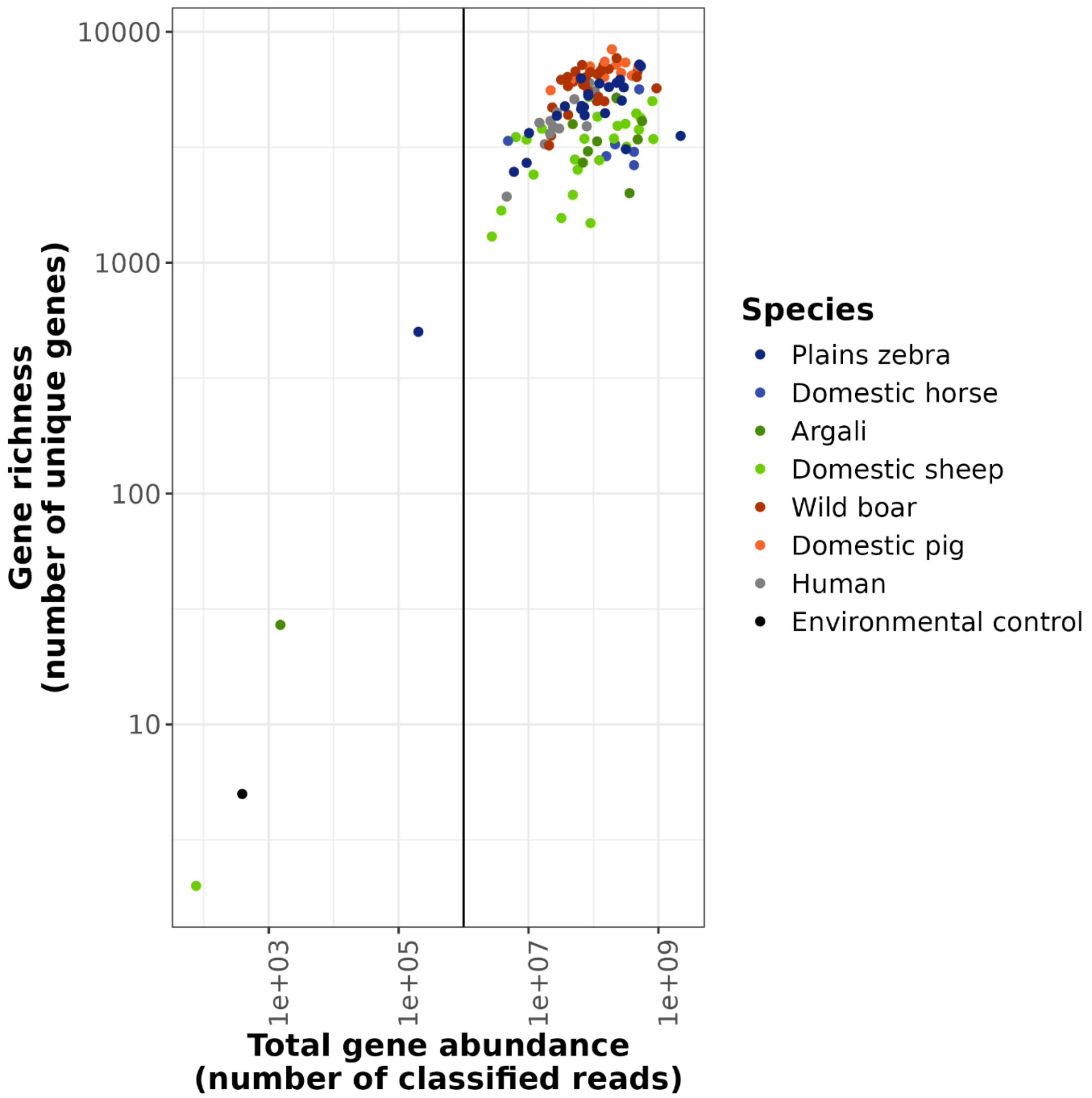


**Supplementary Figure 1.** Number of unique genes plotted against the total gene abundance (number of classified reads) that were retained after decontamination of the gene table. Each point represents a sample and is coloured by the host species. The vertical line at $10^6$ classified reads represents the minimum threshold. Three samples that fell below the threshold were excluded from downstream analyses.

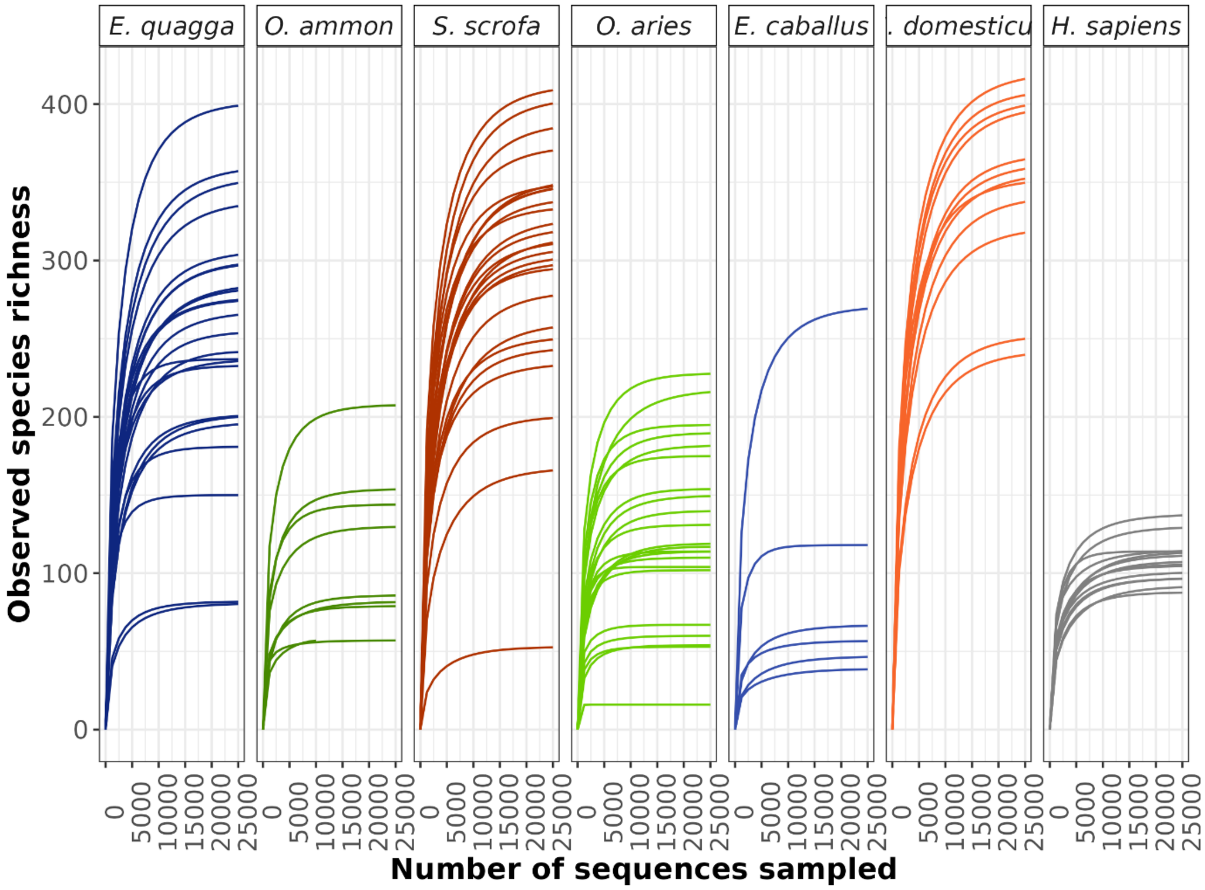


**Supplementary Figure 2.** Rarefaction curves showing estimated species richness at various subsampling depths, from 0 to 25,000 reads. Each panel represents one host species.

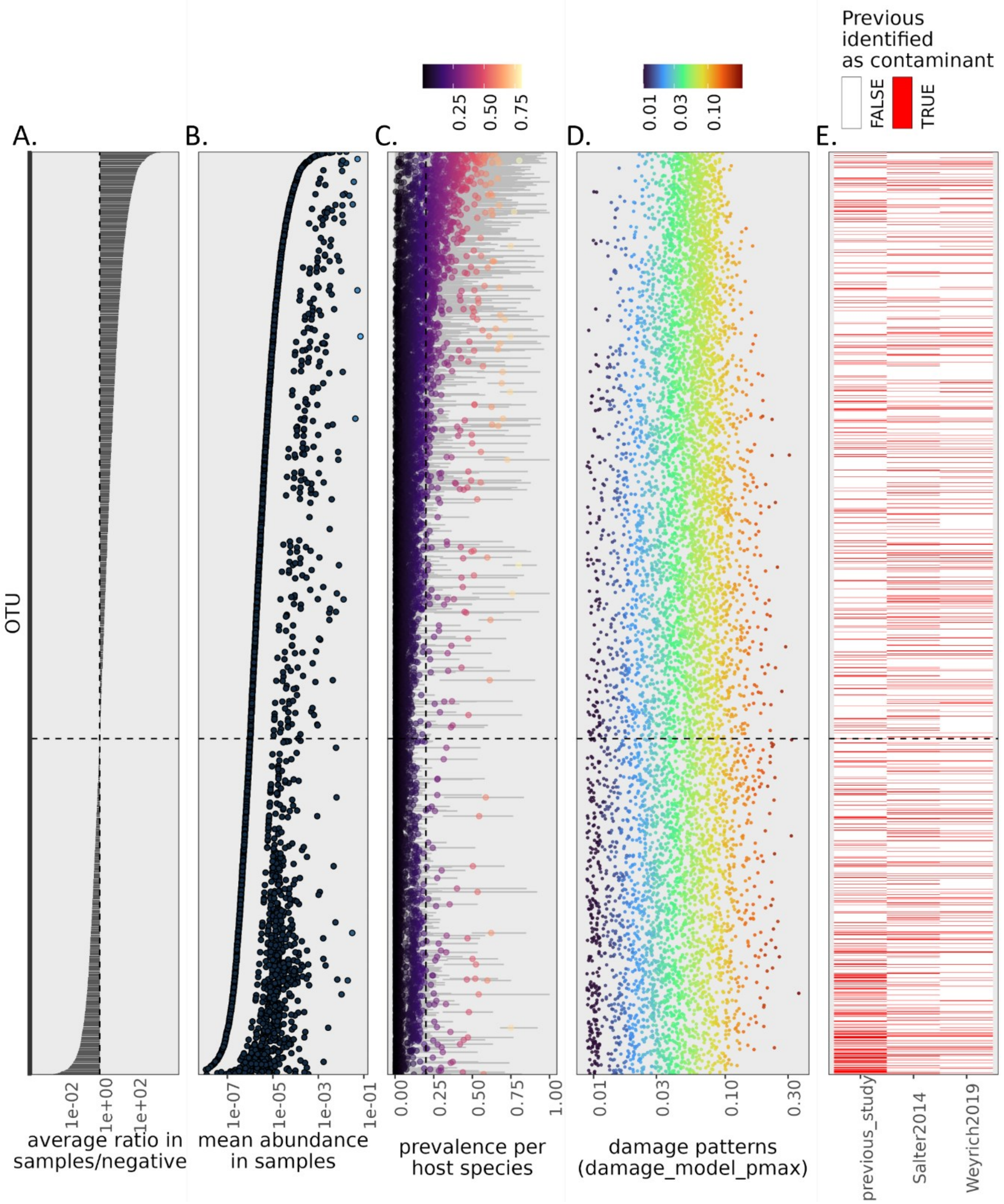


**Supplementary Figure 3**. Dataset decontamination. The y-axis represents all 7,779 microbial species present in dental calculus samples. **A)** The sample/negative control average relative abundance ratio, calculated as the ratio of weighted average relative abundance in samples (see panel B) to the average relative abundance in controls (N=7769). Note log10-transformed x-axis. The vertical dashed line signifies an average relative abundance ratio value of 1, below which taxa were removed as contaminants **B)** Weighted average abundance of each taxon in dental calculus samples (first averaged within host species, and then across host species, to account for differences in sample sizes). **C)** Prevalence of taxa in each host species using a minimum threshold relative abundance of 0.01%. Points indicate the average prevalence across all host species, and are coloured accordingly (see scale bar above the plot), and the grey error bars show the range of values. The vertical dashed line shows the 20% minimum prevalence threshold used for filtering. **D)** Magnitude of post-mortem DNA damage at the 5' end of the read, specifically the "damage_model_pmax" parameter estimated by the PyDamage software (note log10 transformed x-axis). Points indicate the median value for that taxon and are coloured accordingly (see scale bar above the plot). **E)** Information on likely contaminants from the literature. Red bands indicate that a taxon was identified as a contaminant in literature. The first column refers to whether this taxon was identified as a contaminant in Moraitou et al. (2026). The two last columns refer to two published lists of contaminants by Salter et al. (2014) and Weyrich et al. (2021), which were, however, identifying taxa at the genus level. As a result, every species belonging to these contaminant genera has been highlighted in red in the last two columns.

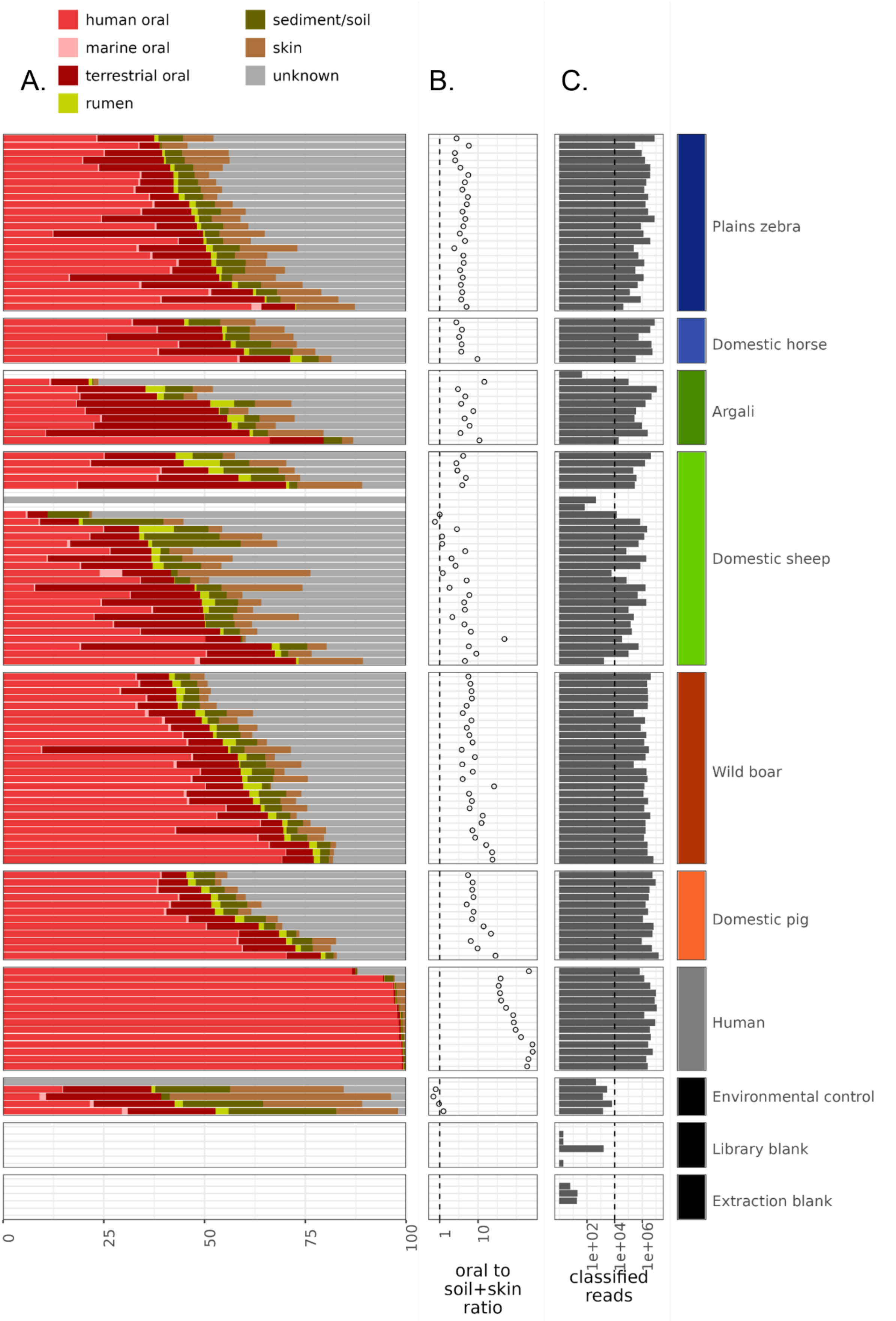


**Supplementary Figure 4**. Assessing sample preservation. **A)** Source-tracking results showing proportions of human oral (medium red), marine mammal oral (pink), terrestrial mammal oral (dark red), rumen (light green), sediment/soil (dark green) and skin (beige) microbial communities, estimated with decOM using a custom source matrix. Note some samples failed to be analysed by this tool and are shown in white. All of these were removed due to an insufficient number of reads (see panel C). **B)** The ratio of the oral proportion (sum of ancient oral and modern oral) to the proportion of skin and soil/sediment (which are the most likely contamination sources) from panel A. Samples with a ratio lower than 1 (those left of the parity line) were consider too poorly preserved and excluded from downstream analyses. **C)** Number of reads classified to the species level. The dashed line shows the threshold of 10,000 classified reads, below which a sample was removed.

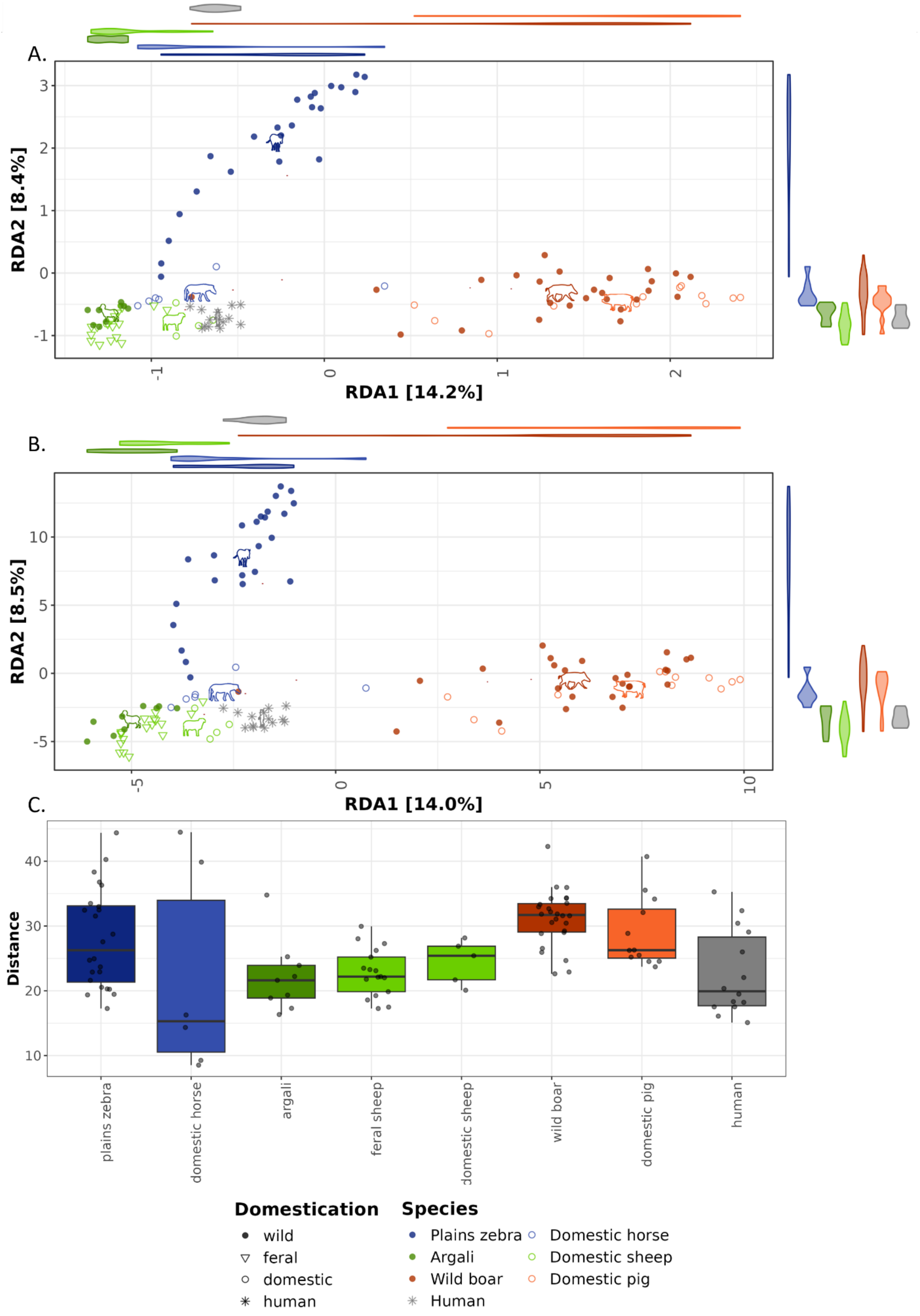


**Supplementary Figure 5. A)** and **B)** Redundancy analysis ordination on presence-absence data **(A)** and PhILR transformed abundances (phylogenetically-informed abundances) **(B)**, using host species as constraints. Host species is shown by different colours, whereas the domestication status is shown with different shapes. The violin plots along the axes show the distributions of the host species along that axis. The phylopic icons indicate the centroid of each host species. **C)** Multivariate variance (calculated as average Aitchison distance of each sample to the group mean) across host groups. Note that feral and domestic sheep (*Ovis aries*) are presented separately.

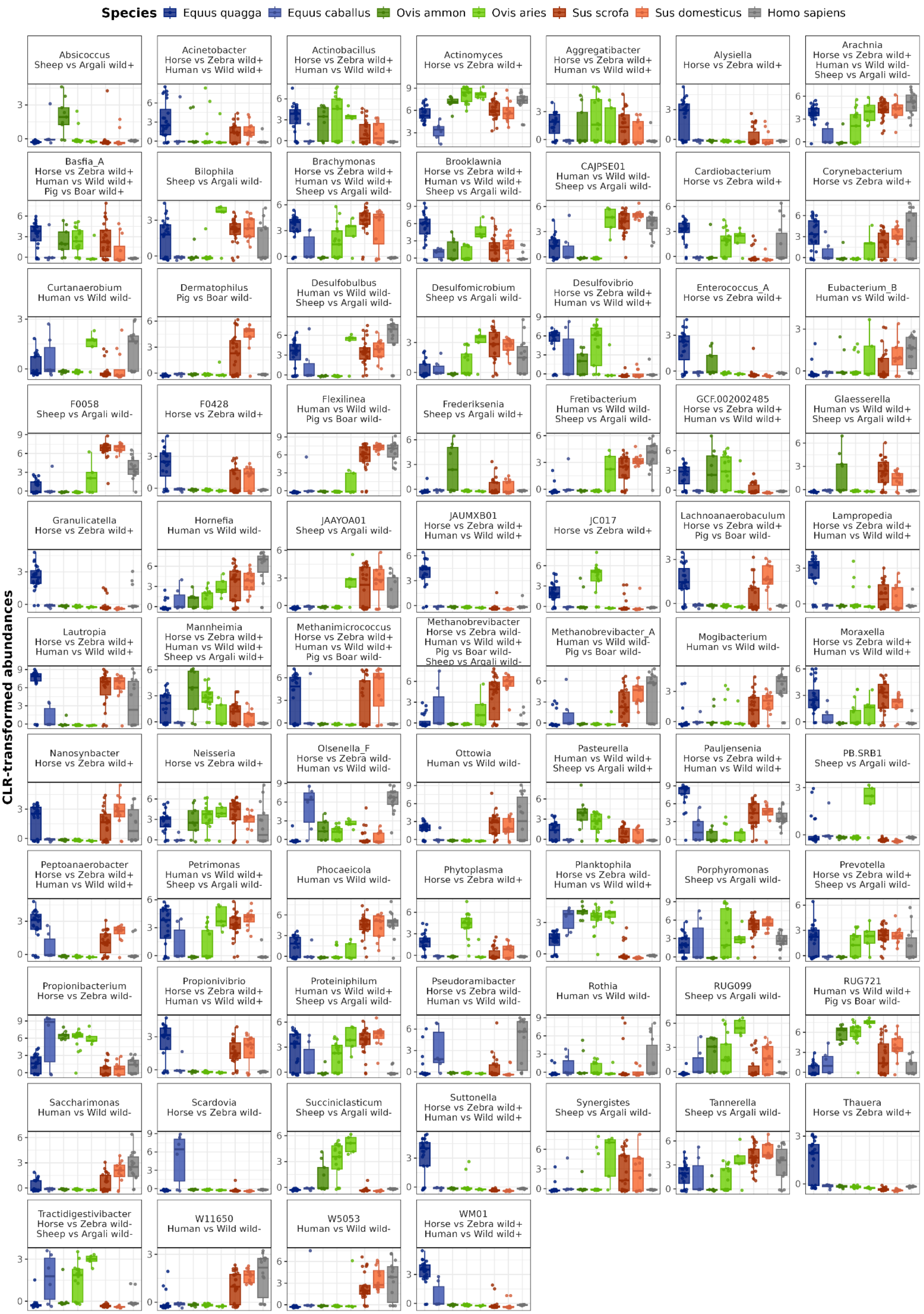
Species
Equus quagga
Equus caballus
Ovis ammon
Ovis aries
Sus scrofa
Sus domesticus
Homo sapiens
CLR-transformed abundances
Absicoccus
Sheep vs Argali wild+
Acinetobacter
Horse vs Zebra wild+
Human vs Wild wild+
Actinobacillus
Horse vs Zebra wild+
Human vs Wild wild+
Actinomyces
Horse vs Zebra wild+
Aggregatibacter
Horse vs Zebra wild+
Human vs Wild wild+
Alysiella
Horse vs Zebra wild+
Arachnia
Horse vs Zebra wild+
Human vs Wild wild-
Sheep vs Argali wild-
Basfia_A
Horse vs Zebra wild+
Human vs Wild wild+
Pig vs Boar wild+
Bilophila
Sheep vs Argali wild-
Brachymonas
Horse vs Zebra wild+
Human vs Wild wild+
Sheep vs Argali wild-
Brooklawnia
Horse vs Zebra wild+
Human vs Wild wild+
Sheep vs Argali wild-
CAJPSE01
Human vs Wild wild-
Sheep vs Argali wild-
Cardiobacterium
Horse vs Zebra wild+
Corynebacterium
Horse vs Zebra wild+
Curtanaerobium
Human vs Wild wild-
Dermatophilus
Pig vs Boar wild-
Desulfobulbus
Human vs Wild wild-
Sheep vs Argali wild-
Desulfomicrobium
Sheep vs Argali wild-
Desulfovibrio
Horse vs Zebra wild+
Human vs Wild wild+
Enterococcus_A
Horse vs Zebra wild+
Eubacterium_B
Human vs Wild wild-
F0058
Sheep vs Argali wild-
F0428
Horse vs Zebra wild+
Flexilinea
Human vs Wild wild-
Pig vs Boar wild-
Frederiksenia
Sheep vs Argali wild+
Fretibacterium
Human vs Wild wild-
Sheep vs Argali wild-
GCF.002002485
Horse vs Zebra wild+
Human vs Wild wild+
Glaesserella
Human vs Wild wild+
Sheep vs Argali wild+
Granulicatella
Horse vs Zebra wild+
Hornefia
Human vs Wild wild-
JAAYOA01
Sheep vs Argali wild-
JAUMXB01
Horse vs Zebra wild+
Human vs Wild wild+
JC017
Horse vs Zebra wild+
Lachnoanaerobaculum
Horse vs Zebra wild+
Pig vs Boar wild-
Lampropedia
Horse vs Zebra wild+
Human vs Wild wild+
Lautropia
Horse vs Zebra wild+
Human vs Wild wild+
Mannheimia
Horse vs Zebra wild+
Human vs Wild wild+
Sheep vs Argali wild+
Methanimicrococcus
Horse vs Zebra wild+
Human vs Wild wild+
Pig vs Boar wild-
Methanobrevibacter
Horse vs Zebra wild-
Human vs Wild wild+
Pig vs Boar wild-
Sheep vs Argali wild-
Methanobrevibacter_A
Human vs Wild wild-
Pig vs Boar wild-
Mogibacterium
Human vs Wild wild-
Moraxella
Horse vs Zebra wild+
Human vs Wild wild+
Nanosynbacter
Horse vs Zebra wild+
Neisseria
Horse vs Zebra wild+
Olsenella_F
Horse vs Zebra wild-
Human vs Wild wild-
Ottowia
Human vs Wild wild-
Pasteurella
Human vs Wild wild+
Sheep vs Argali wild+
Pauljensenia
Horse vs Zebra wild+
Human vs Wild wild+
PB.SRB1
Sheep vs Argali wild-
Peptoanaerobacter
Horse vs Zebra wild+
Human vs Wild wild+
Petrimonas
Human vs Wild wild+
Sheep vs Argali wild-
Phocaeicola
Human vs Wild wild-
Phytoplasma
Horse vs Zebra wild+
Planktophila
Horse vs Zebra wild-
Human vs Wild wild+
Porphyromonas
Sheep vs Argali wild-
Prevotella
Horse vs Zebra wild+
Sheep vs Argali wild-
Propionibacterium
Horse vs Zebra wild-
Propionivibrio
Horse vs Zebra wild+
Human vs Wild wild+
Proteiniphilum
Human vs Wild wild+
Sheep vs Argali wild-
Pseudoramibacter
Horse vs Zebra wild-
Human vs Wild wild-
Rothia
Human vs Wild wild-
RUG099
Sheep vs Argali wild-
RUG721
Human vs Wild wild+
Pig vs Boar wild-
Saccharimonas
Human vs Wild wild-
Scardovia
Horse vs Zebra wild-
Succiniclasticum
Sheep vs Argali wild-
Suttonella
Horse vs Zebra wild+
Human vs Wild wild+
Synergistes
Sheep vs Argali wild-
Tannerella
Sheep vs Argali wild-
Thauera
Horse vs Zebra wild+
Tractidigestivibacter
Horse vs Zebra wild-
Sheep vs Argali wild-
W11650
Human vs Wild wild-
W5053
Human vs Wild wild-
WM01
Horse vs Zebra wild+
Human vs Wild wild+

**Supplementary Figure 6. Abundances of differentially abundant taxa across host groups.** CLR-transformed abundances of genera that were identified as differentially abundant (MCMCglmm pMCMC < 0.01) in any of the three within-genus comparisons: within *Equus* (domestic horse against plains zebra), within *Ovis* (domestic sheep against argali) and within *Sus* (domestic pig versus wild boar). To put these results into context, we are also noting if these taxa were enriched or depleted among wild animals in comparison to humans. The type of significant relationship ('wild+' for enriched and 'wild-' for depleted in wild animals) and the specific host taxon comparison are indicated under the bacterial genus name. Feral sheep abundances are plotted but not considered in the differential abundance analyses. The taxa are presented in alphabetical order.

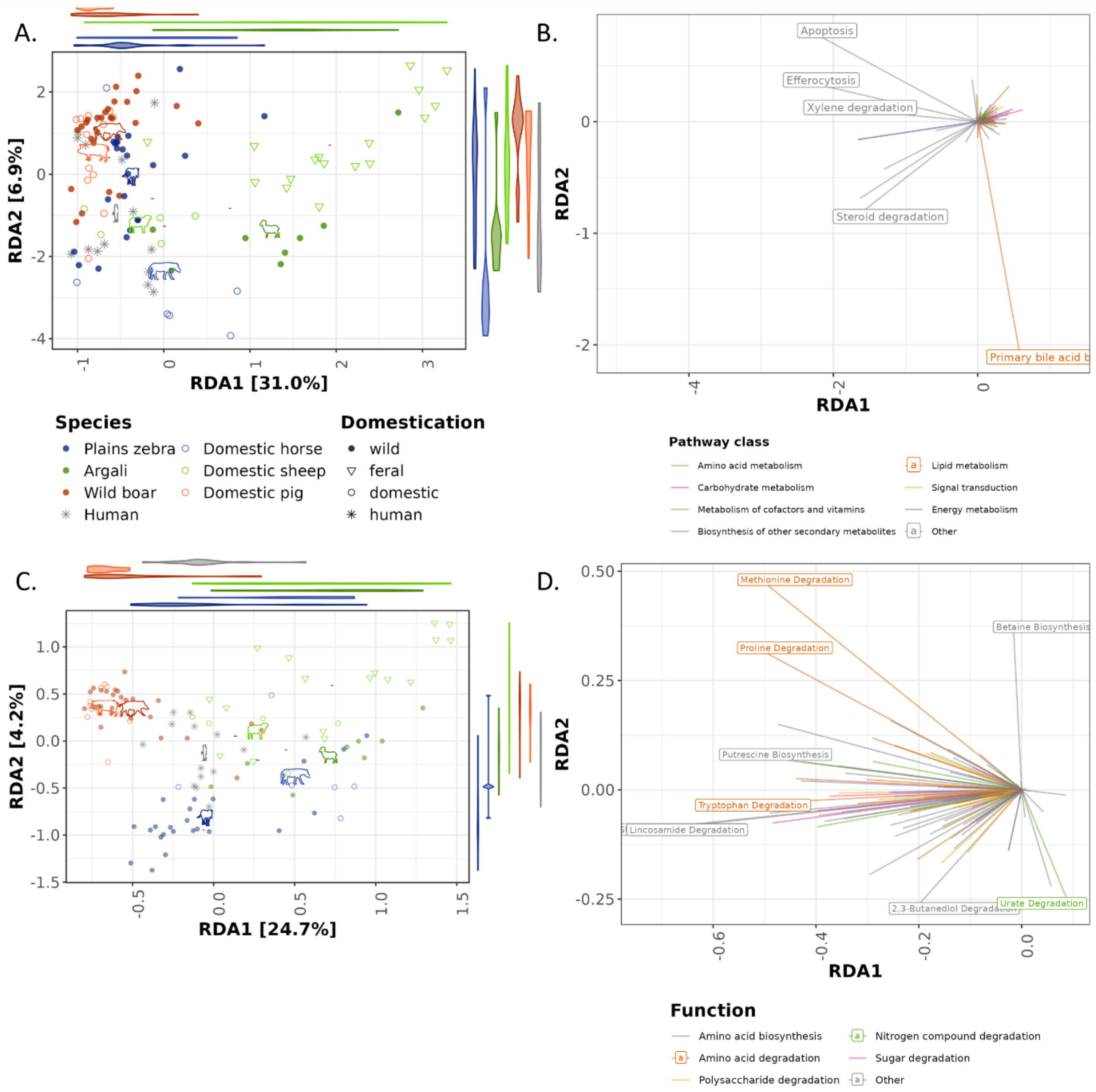


**Supplementary Figure 7. Left column:** Redundancy analysis ordinations on functional profiles using host species as constraints. Host species are shown by different colours, whereas the domestication status is shown with different shapes. The violin plots along the axes show the distributions of the host species along that axis. The phylopic icons indicate the centroid of each host species. **Right column:** Arrows indicating the contributions of variables to the ordination plots in the left column**.** **A)** Redundancy analysis ordination on CLR-transformed abundances of KEGG pathways. **B)** Contributions of KEGG pathways to the variation in ordination (A). The direction of each arrow indicates the gradient along which the pathway increases in abundance, and the length is proportional to the strength of the contribution to the ordination axes. The colour corresponds to the KEGG pathway class (with less common classes grouped as "Other"). Labels are shown for the five pathways with the strongest contributions. **C)** Redundancy analysis ordination on the completeness of functional traits as identified by distillR. **D)** Contributions of functional traits pathways to the variation in ordination (C). The direction of each arrow indicates the gradient along which the functional trait increases in completeness, and the length is proportional to the strength of the contribution to the ordination axes. The colour corresponds to the function grouping as defined by distillR (with less common functions grouped as "Other"). Labels are shown for the five pathways with the strongest contributions.

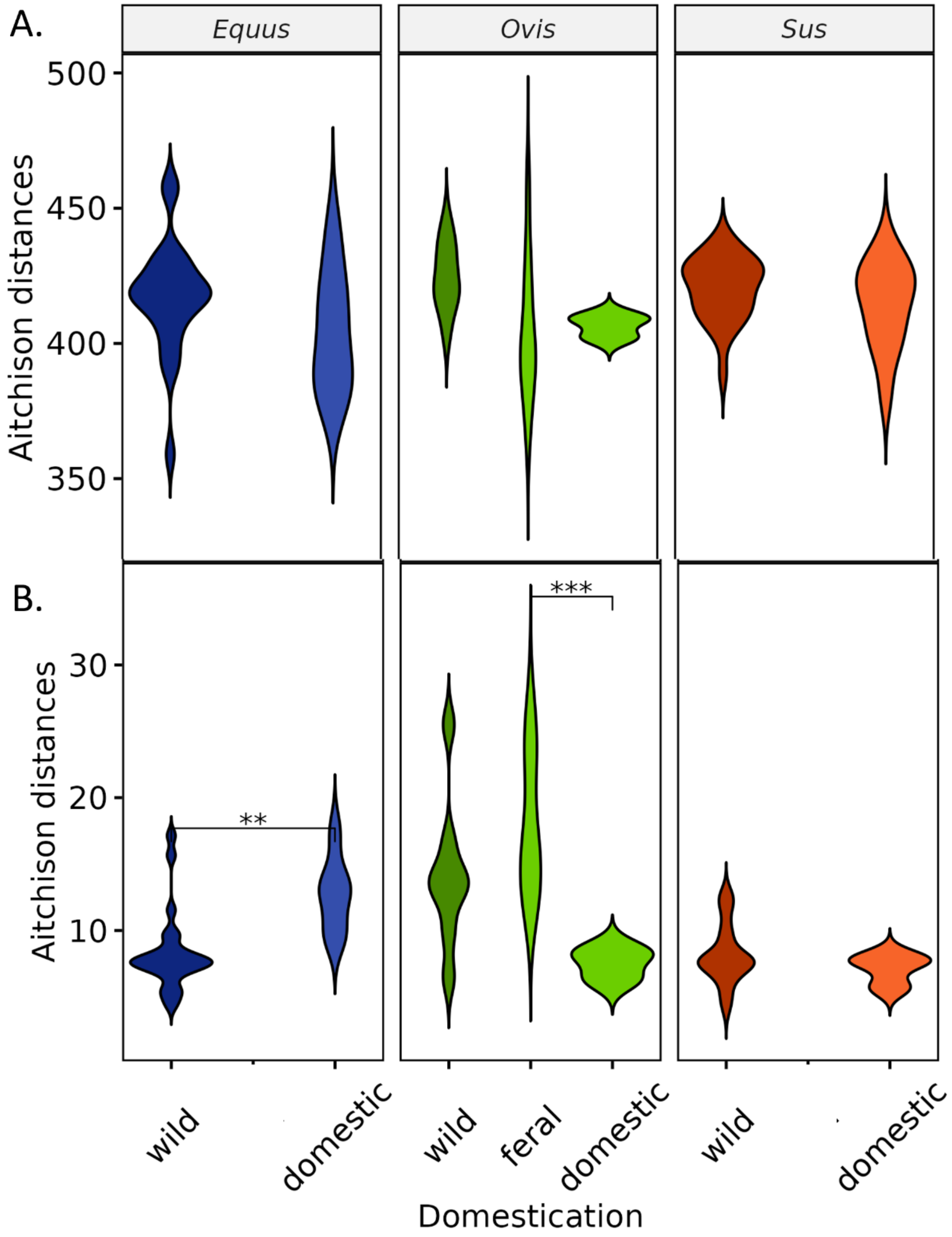


**Supplementary Figure 8.** Distance of the functional profiles of each sample to human centroid. **A)** Aitchison distances on gene abundances. **B)** Aitchison distances on KEGG pathway abundances. Asterisks indicate significant differences between the different domestication categories of a host genus, identified using a Kruskal-Wallis test ($p < 0.05$).

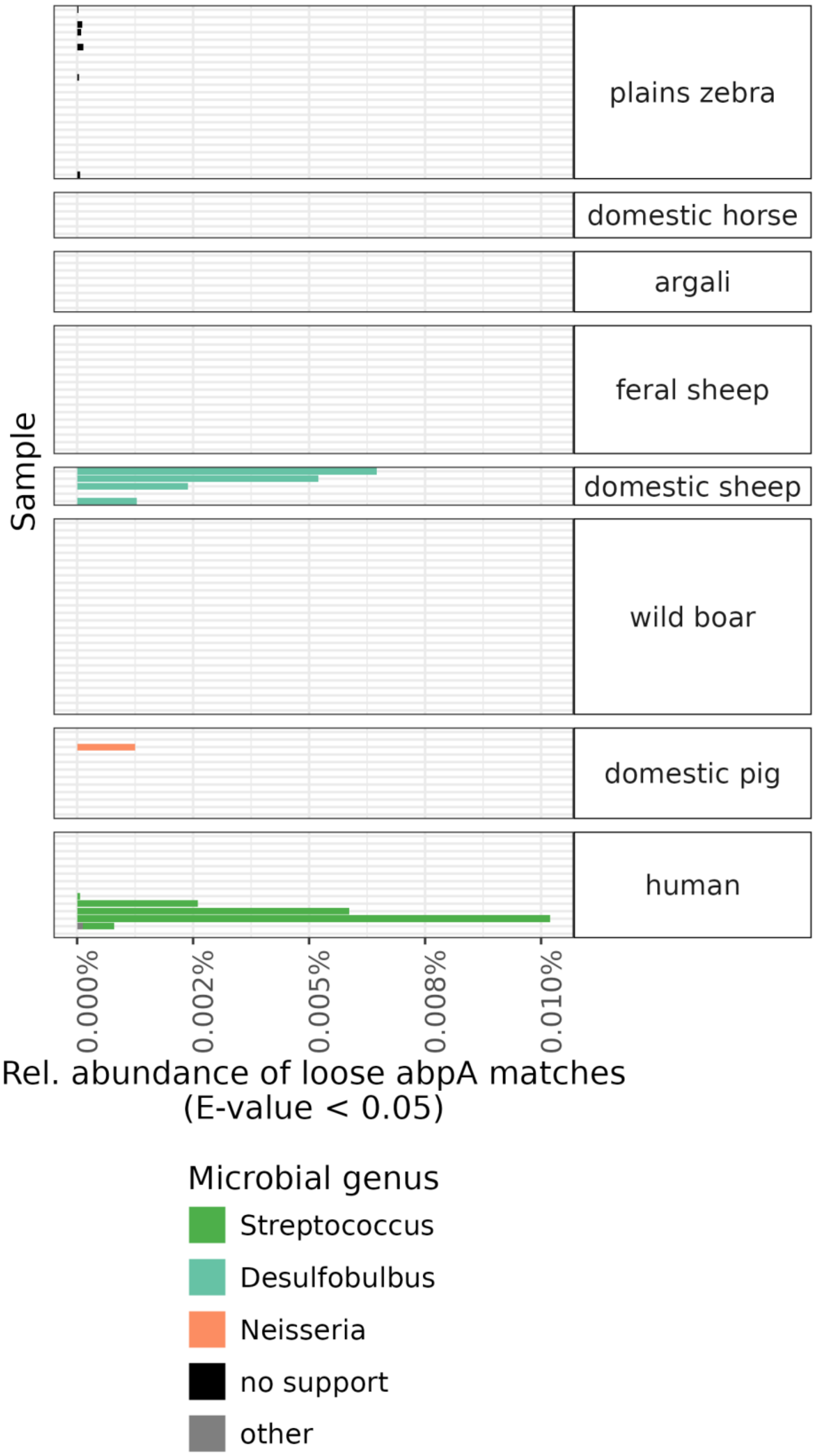


**Supplementary Figure 9.** Relative abundance of putative genes matching to abpA amylase-binding protein with E-value less than 0.05. The bar colours show the microbial genus containing the hit, with "no support" indicating hits to contigs that could not be resolved at the bacterial genus level.

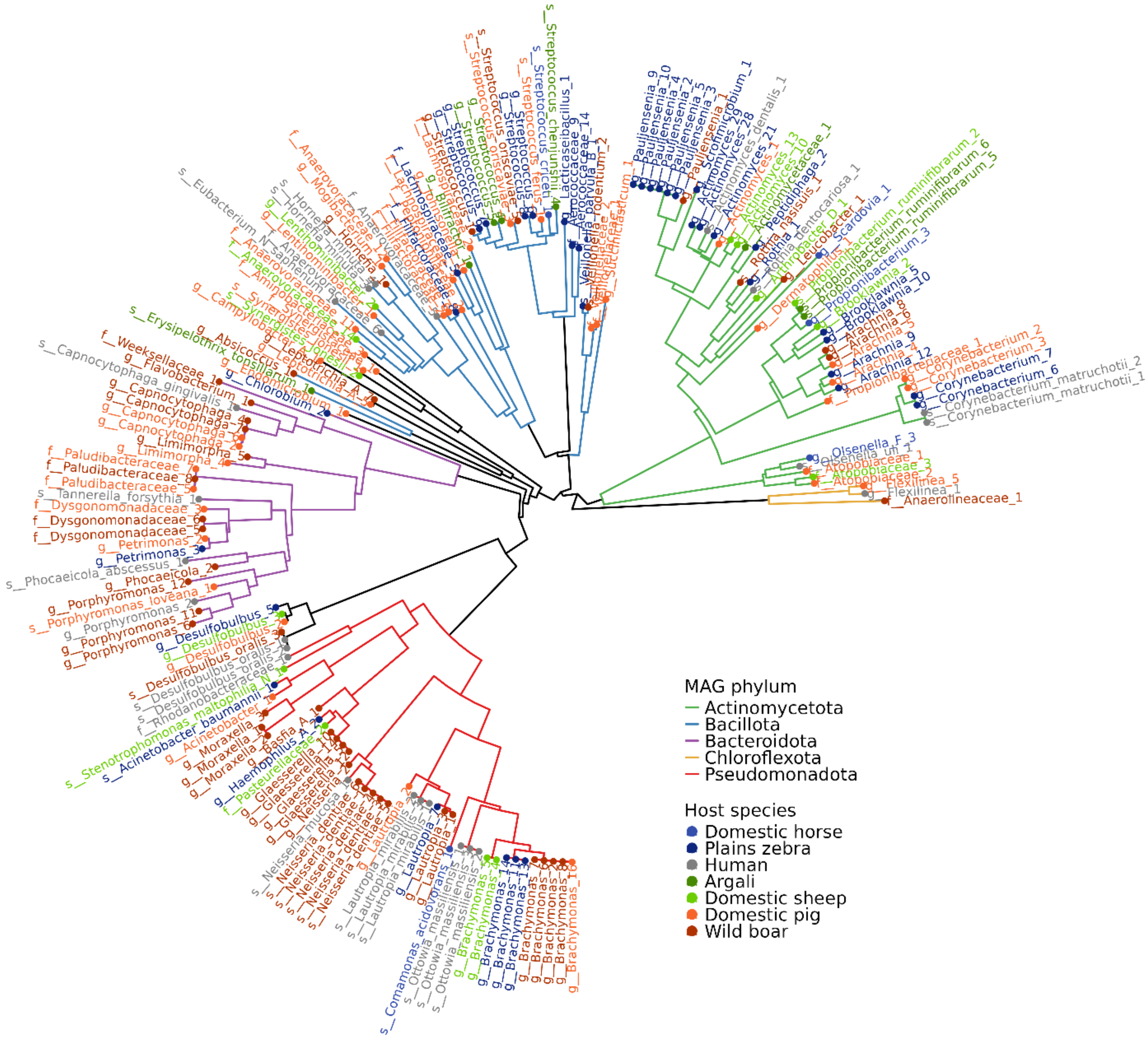


**Supplementary Figure 10.** Bacterial phylogeny created using the GTDB-tk ‘de_novo_wf’ workflow, for bacterial MAGs with completeness >75% and contamination <5%. Tree branches are coloured according to the phylum to which the MAG was assigned. The tips and labels are coloured according to the host species from which the MAG was assembled.

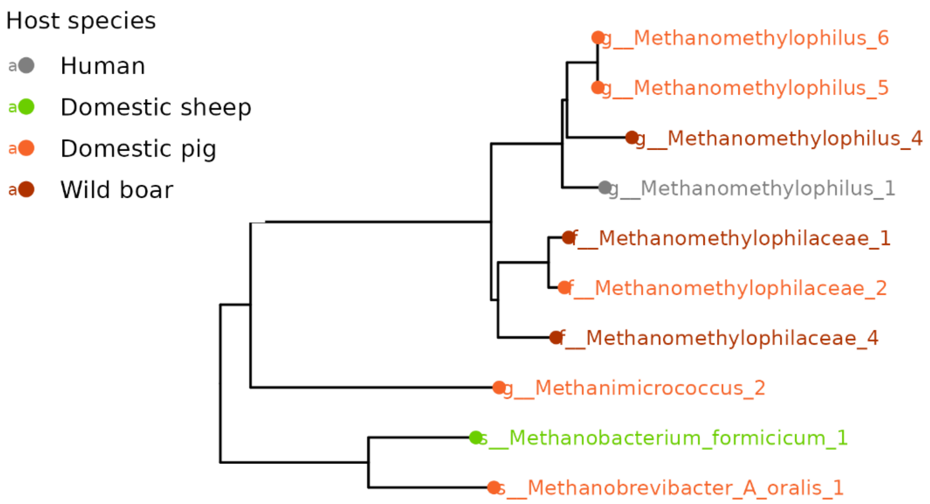


**Supplementary Figure 11.** Archaeal phylogeny created using the GTDB-tk 'de_novo_wf' workflow for archaeal MAGs with completeness >75% and contamination <5%. The tips and labels are coloured according to the host species from which the MAG was assembled.

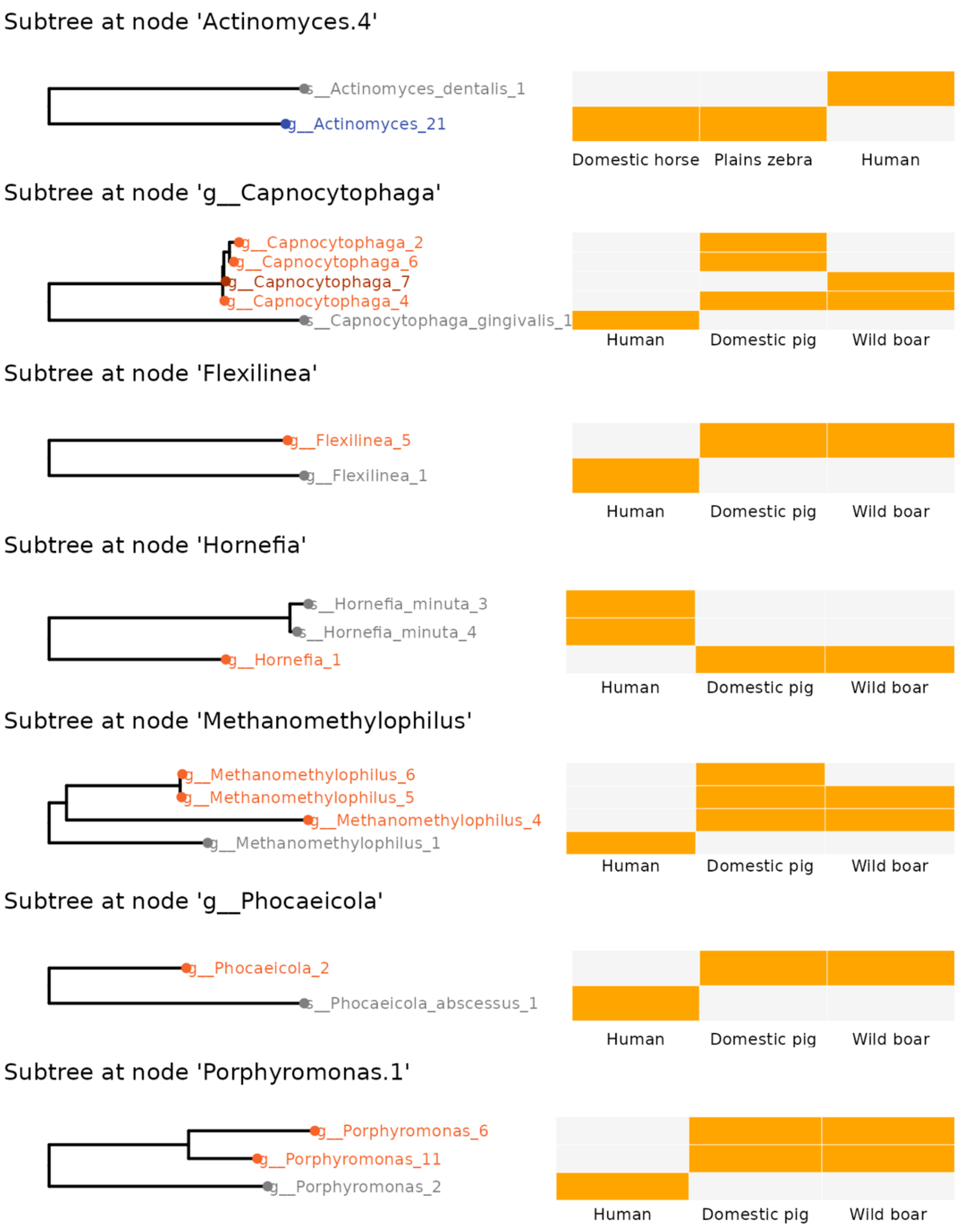


**Supplementary Figure 12. Phylogenetic subtrees of the nodes that were found to follow a ((D, W), H) topological pattern.** In six out of seven subtrees, this is more specifically a ((domestic pig, wild boar), human) pattern. Heatmaps indicate MAG presence across the three host species (grey = absent, orange = present). The colour of the points and labels of the phylogenetic trees indicate the host species in which the MAGs were assembled (grey = human, orange = domestic pig, dark orange = wild boar, blue = plains zebra). All are nodes of the bacterial phylogeny, except for *Methanomethylophilus,* which is of the archaeal phylogeny.
